# Active Enterohepatic Cycling is Not Required for the Choleretic Actions of 24-*nor*Ursodeoxycholic Acid in Mice

**DOI:** 10.1101/2021.03.06.430841

**Authors:** Jianing Li, Jennifer K. Truong, Kimberly Pachura, Anuradha Rao, Sanjeev Gambeer, Claudia Daniela Fuchs, Saul J. Karpen, Michael Trauner, Paul A. Dawson

**Author notes:** JL and JKT contributed equally to this work. Address correspondence to: Paul A. Dawson, Ph.D., Pediatric Gastroenterology, Hepatology and Nutrition, Emory University School of Medicine, Health Sciences Research Building, 1760 Haygood Drive, NE, Suite 200E, Atlanta, Georgia 30322.

## Abstract

The superior ability of *nor*ursodeoxycholic acid (*nor*UDCA) to induce a bicarbonate-rich hypercholeresis has been attributed to its ability to undergo cholehepatic shunting and *nor*UDCA is currently being evaluated as a therapeutic for forms of liver disease. The goal of this study was to use mouse models to investigate contributions of bile acid transporters to the choleretic actions of *nor*UDCA. Here, we show that the apical sodium-dependent bile acid transporter (ASBT) and Organic solute transporter-alpha (OSTα) are dispensable for *nor*UDCA-stimulation of bile flow and biliary bicarbonate secretion in mice. Analysis of the liver transcriptome revealed that *nor*UDCA induced hepatic expression of a limited number of transporter genes, particularly organic anion transporting polypeptide 1a4 (Oatp1a4). However, *nor*UDCA potently stimulated a bicarbonate-rich hypercholeresis in Oatp1a/1b-deficient mice. Blocking intestinal bile acid reabsorption by co-administration of an ASBT inhibitor or bile acid sequestrant did not impact the ability of *nor*UDCA to induce bile flow in wildtype mice. The results support the concept that these major bile acid transporters are not directly involved in the absorption, cholehepatic shunting, or choleretic actions of *nor*UDCA. Additionally, the findings support further investigation of the therapeutic synergy between *nor*UDCA and ASBT inhibitors or bile acid sequestrants for cholestatic liver disease.

## Introduction

*nor*Ursodeoxycholic acid (*nor*UDCA) is a synthetic C-23 side chain-shortened analog of the hydrophilic native bile acid ursodeoxycholic acid (UDCA) and is resistant to side-chain conjugation with glycine or taurine (1). The pharmacological properties and physiological actions of *nor*UDCA make it a therapeutic candidate for a variety of cholestatic liver diseases (1). In preclinical studies, oral administration of *nor*UDCA reduced liver injury and biliary fibrosis in bile duct ligated mice and in *Abcb4/Mdr2^−/−^* mice, whereas administration of UDCA aggravated liver and bile duct injury (2, 3). In those models, *nor*UDCA induced detoxification and renal elimination of bile acids and exhibited anti-proliferative, anti-fibrotic, and anti-inflammatory properties (2–6). In Phase 2 clinical trials, administration of *nor*UDCA for 12 weeks reduced serum alkaline phosphatase (ALP) and other liver enzyme markers of cholestasis in patients with Primary Sclerosing Cholangitis (PSC) (7), and reduced serum alanine aminotransferase (ALT) in patients with non-alcoholic fatty liver disease (8).

Administration of side chain-shortened dihydroxy bile acids such as *nor*UDCA produce a bicarbonate-rich bile flow in excess of what can be explained by their osmotic effects (1). The superior ability of *nor*UDCA versus UDCA to induce a hypercholeresis has been attributed to *nor*UDCA’s ability to evade side chain conjugation (amidation) to glycine or taurine and undergo cholehepatic shunting. In the pathway proposed by Hofmann (1, 9), unconjugated *nor*UDCA is secreted by hepatocytes into bile and absorbed in protonated form by cholangiocytes lining the biliary tract, thereby generating a bicarbonate ion from biliary CO_2_. *nor*UDCA then crosses the biliary epithelium and enters the periductular capillary plexus, which drains into the portal vein (or directly into the hepatic sinusoids), delivering *nor*UDCA for uptake by hepatocytes and subsequent resecretion into bile. Unmodified *nor*UDCA that escapes absorption in the biliary tract travels along with other biliary constituents into the small intestine where it is reabsorbed and carried in the enterohepatic circulation back to the liver for uptake and resecretion into bile. In this fashion, *nor*UDCA engages in multiple rounds of cholehepatic shunting or a combination of enterohepatic and cholehepatic cycling before being converted to a more polar metabolite by hepatic phase 1 or phase 2 metabolism (primarily phase 2 glucuronidation) and eliminated from the body in the urine or feces (1, 9–11). The physicochemical and permeability properties of *nor*UDCA (Supplemental Table 1) are generally consistent with a role for passive diffusion in the absorption of *nor*UDCA (12). However, carrier-mediated cellular uptake and export mechanisms also play prominent roles in the uptake and disposition of many drugs and endobiotics (13–15), and the contribution of bile acid and organic anion transporters to the absorption and cholehepatic shunting of *nor*UDCA has not been fully explored (16). In this study, we have taken a genetic approach to determine whether the major bile acid transporters ASBT and OSTα-OSTβ and an active enterohepatic circulation are required for the hypercholeretic actions of *nor*UDCA.

## Results

To determine if the major bile acid transporter Asbt and an active enterohepatic circulation of bile acids are required for the bicarbonate-rich choleresis induced by *nor*UDCA, we examined bile flow and biliary bicarbonate output in background strain-matched WT and *Asbt^−/−^* mice fed chow or chow plus 0.5% *nor*UDCA for 7 days. The experimental scheme and morphological response to *nor*UDCA administration is shown in Supplemental Figure 2. Administration of *nor*UDCA to WT and *Asbt^−/−^* mice for 7 days tended to reduce body weight (Supplemental Figure 2B, C), but did not affect small intestinal length or weight, colon length or weight, or kidney weight (data not shown). The liver weight and liver to body weight ratio were increased in both genotypes with *nor*UDCA treatment (Supplemental Figure 2D, E). However, analysis of H&E-stained liver sections revealed no apparent histological differences between the genotypes or treatment groups (Supplemental Figure 2F), and plasma chemistries were not significantly different between the chow and *nor*UDCA-fed groups for both genotypes (Supplemental Table 2).

The effect of *nor*UDCA-administration on bile flow and biliary solute output is shown in Figure 1 and summarized in Table 1. On the rodent chow diet, bile flow, bicarbonate concentration, biliary bicarbonate output, and bile pH were similar in WT and *Asbt^−/−^* mice. In agreement with a block in ileal active reabsorption of bile acids, the concentration and biliary output of bile acids was reduced by more than 50% in chow-fed *Asbt^−/−^* versus WT mice (Figure 1D, E). As compared to chow-fed mice, administration of *nor*UDCA increased the bile flow rate by 5 to 6-fold, biliary bicarbonate concentration by 2-fold, and bicarbonate output more than 10-fold in both WT and *Asbt^−/−^* mice (Figures 1A, B, C; Table 1). *nor*UDCA-feeding also increased bile acid output by approximately 4-fold and 8-fold in WT and *Asbt^−/−^* mice, respectively (Figures 1D, E). Since the ability of *nor*UDCA to stimulate a bicarbonate-rich choleresis is thought to be secondary to its potential for cholehepatic shunting and enrichment in bile, biliary bile acid composition was determined for chow and *nor*UDCA-treated WT and *Asbt^−/−^* mice. The output and relative proportion of each bile acid species are shown (Figures 1E, 1F). As compared to chow-fed WT mice, *Asbt^−/−^* mice had a more hydrophobic bile acid composition, with reduced relative amounts of 6-hydroxylated bile acid species such as tauro-β-muricholic acid (TβMCA), and increased amounts of taurocholic acid (TCA) and its gut microbiota-derived product taurodeoxycholic acid (TDCA). Following administration of *nor*UDCA, the biliary bile acid composition became more hydrophilic in *Asbt*^−/−^ mice and remarkably similar to WT mice, with *nor*UDCA accounting for approximately 60% of the total biliary bile acids in both genotypes. There was also a large reduction in the proportion of TCA and TDCA in *Asbt^−/−^* mice following *nor*UDCA treatment. The biliary bile acid hydrophobicity changes are reflected in the calculated Hydrophobicity Index, which decreased from +0.166 to −0.483 in *Asbt^−/−^* mice with *nor*UDCA feeding but was largely unchanged in WT mice (calculated Hydrophobicity Index value of −0.453 versus −0.489 in chow and *nor*UDCA-fed mice, respectively). For comparison, the amounts of different bile acid species excreted into the feces are shown (Supplemental Figure 3). Under chow-fed conditions, the fecal bile acid content was approximately 5-fold greater in *Asbt^−/−^* versus WT mice and included a higher proportion of cholic acid and deoxycholic acid. Administration of *nor*UDCA in the diet increased the fecal bile acid content in both WT and *Asbt^−/−^* mice and shifted the endogenous bile acid composition toward the 6-hydroxylated muricholate species. The increase in fecal bile acid levels was driven primarily by the exogenous *nor*UDCA, however the amount of endogenous bile acid in feces was also increased in both WT mice and *Asbt^−/−^* mice after administration of *nor*UDCA.

**Table 1.** Bile Flow and Composition in WT and *Asbt^−/−^* Mice Fed Chow or *nor*UDCA Diet. Values are expressed as the mean ± SD. The number of mice per group are indicted (n). Values with different superscript letters are significantly different (*P* < 0.05) according to ordinary two-way ANOVA and Sidak’s multiple comparisons test. BLD, below level of detection; LW, liver weight.

| Variable | Chow |  | *nor*UDCA |  |
| --- | --- | --- | --- | --- |
|  | WT (6) | *Asbt*<sup>-/-</sup> (7) | WT (7) | *Asbt*<sup>-/-</sup> (7) |
| Bile Flow (μl/g LW/min) | 1.1 ± 0.1<sup>a</sup> | 0.9 ± 0.2<sup>a</sup> | 5.4 ± 0.6<sup>b</sup> | 6.0 ± 2.5<sup>b</sup> |
| Bicarbonate (mM) | 32 ± 4<sup>a</sup> | 30 ± 4<sup>a</sup> | 62 ± 5<sup>b</sup> | 54 ± 6<sup>c</sup> |
| Bicarbonate Output (nmol/g LW/min) | 34 ± 7<sup>a</sup> | 27 ± 10<sup>a</sup> | 335 ± 58<sup>b</sup> | 324 ± 117<sup>b</sup> |
| Bile pH | 7.6 ± 0.1<sup>a</sup> | 7.7 ± 0.1<sup>a</sup> | 7.9 ± 0.1<sup>b</sup> | 7.7 ± 0.1<sup>a</sup> |
| Bile Acid (mM) | 26.8 ± 5.8<sup>a</sup> | 12.8 ± 2.5<sup>b</sup> | 22.5 ± 8.4<sup>a,b</sup> | 16.5 ± 7.2<sup>a,b</sup> |
| Bile Acid Output (nmol/g LW/min) | 28.9 ± 8.8<sup>a</sup> | 11.8 ± 5.4<sup>a</sup> | 123.6 ± 52.2<sup>b</sup> | 96.2 ± 45.8<sup>b</sup> |
| Glutathione (mM) | 0.65 ± 0.81<sup>a</sup> | 1.41 ± 0.82<sup>a</sup> | 0.74 ± 0.37<sup>a</sup> | 0.80 ± 0.42<sup>a</sup> |
| Glutathione Output (nmol/g LW/min) | 0.7 ± 1.0<sup>a</sup> | 1.3 ± 0.9<sup>a</sup> | 4.0 ± 2.1<sup>b</sup> | 4.5 ± 2.5<sup>b</sup> |
| Cholesterol (μg/μl) | 0.17 ± 0.07<sup>a,b</sup> | 0.2 ± 0.06<sup>a</sup> | 0.1 ± 0.04<sup>b</sup> | 0.1 ± 0.07<sup>a,b</sup> |
| Cholesterol Output (μg/g LW/min) | 0.2 ± 0.1<sup>a</sup> | 0.2 ± 0.1<sup>a</sup> | 0.4 ± 0.2<sup>a,b</sup> | 0.6 ± 0.4<sup>b</sup> |
| Phospholipid (mM) | 2.4 ± 1.3 | 3.0 ± 1.4 | BLD | BLD |
| Phospholipid Output (nmol/g LW/min) | 2.7 ± 1.8 | 2.9 ± 1.9 | BLD | BLD |

Values are expressed as the mean ± SD. The number of mice per group are indicated (n). Values with different superscript letters are significantly different ( $P < 0.05$ ) according to ordinary two-way ANOVA and Sidak's multiple comparisons test. BLD, below level of detection; LW, liver weight.

**Figure 1.**
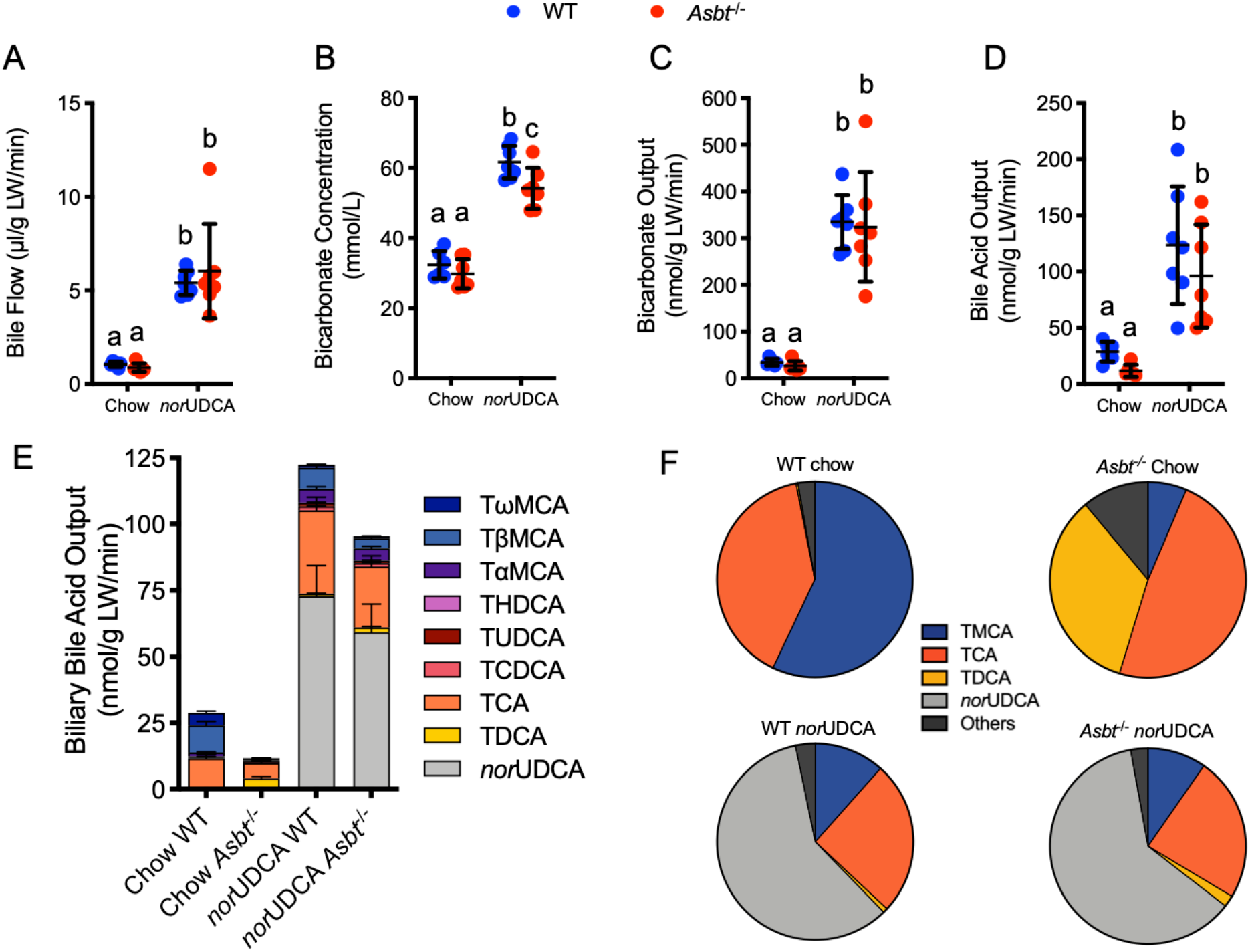
*nor*UDCA treatment increases bile flow and biliary bicarbonate and solute output in WT and *Asbt^−/−^* mice. (**A**) Bile flow. (**B**) Biliary bicarbonate concentration. (**C**) Bicarbonate output. (**D**) Biliary bile acid output. (**E**) Biliary bile acid species output (mean + SEM). (**F**) Biliary bile acid composition expressed as pie charts. Unless indicated, mean + SD are shown; n = 6-7 mice per group. Distinct lowercase letters indicate significant differences between groups (*P* < 0.05).

The effect of *nor*UDCA-feeding on the output of other biliary solutes in WT and *Asbt^−/−^* mice are shown in Table 1. The total glutathione concentration and output tended to be higher in chow-fed *Asbt^−/−^* versus WT mice. This may be a mechanism to increase bile acid-independent bile flow in order to compensate for interruption of the bile acid enterohepatic circulation and reductions in bile acid-dependent bile flow. Administration of *nor*UDCA to WT and *Asbt^−/−^* mice did not change the biliary glutathione concentration, but increased glutathione output by 3 to 4-fold in both genotypes. Biliary cholesterol levels were slightly decreased in WT and *Asbt^−/−^* mice fed the *nor*UDCA diet, but total cholesterol output was increased versus chow-fed mice due to increases in bile flow. In contrast to biliary cholesterol, administration of *nor*UDCA dramatically reduced biliary phospholipid secretion in both WT and *Asbt^−/−^* mice, in agreement with previous studies (1–3) and has been attributed to reduced surface activity and ability of norUDCA to extract phospholipid from the canalicular membrane (10). Overall, these findings suggest that the Asbt is not required for the absorption of *nor*UDCA or its ability to stimulate a bicarbonate-rich hypercholeresis in mice.

OSTα-OSTβ is a heteromeric bi-directional facilitative transporter and is responsible for bile acid export across the basolateral membrane of various epithelium. Similar to the ASBT, OSTα-OSTβ is expressed by ileal enterocytes and cholangiocytes. However, OSTα-OSTβ is also expressed at lower levels in proximal small intestine and in colon, where it may be involved in the export of bile acids that were taken up across the apical membrane by passive diffusion (17, 18). Due to their higher pKa, a fraction of unconjugated and glycine-conjugated bile acids are protonated and gain entry to the epithelium by nonionic diffusion (19). However, once inside the cytoplasmic compartment, weak acids ionize at this neutral pH, potentially impeding their exit from the cell by passive diffusion and necessitating the requirement for an efflux carrier such as OSTα-OSTβ. To determine if OSTα-OSTβ may be contributing to the absorption and bicarbonate-rich choleresis induced by *nor*UDCA, we examined bile flow and biliary bicarbonate output in background strain-matched WT and *Ostα^−/−^Asbt^−/−^* mice fed chow or chow plus 0.5% *nor*UDCA for 7 days. *Ostα^−/−^Asbt^−/−^* mice were selected for these studies in place of *Ostα^−/−^* mice because inactivation of the Asbt protects *Ostα^−/−^* mice from ileal injury and attenuates the associated adaptive changes such as lengthening of the small intestine, and ileal histological alterations such as villous blunting, increased numbers of mucin-producing cells, and increased cell proliferation (20). These changes were predicted to complicate interpretation of the findings with regards to the role of Ostα-Ostβ transport activity in the intestinal absorption and choleretic actions of *nor*UDCA. The experimental scheme and morphological response to *nor*UDCA administration is shown in Supplemental Figure 4. As with the WT and *Asbt^−/−^* mice, liver to body weight ratio was increased in *Ostα^−/−^Asbt^−/−^* and matched WT mice with *nor*UDCA treatment (Supplemental Figure 4E). Compared to chow-fed mice, administration of *nor*UDCA increased bile flow rate by 5 to 6-fold, biliary bicarbonate concentration by 2-fold, bicarbonate output by more than 10-fold, and glutathione output by 5 to 6-fold in WT mice and background strain-matched mice lacking both Asbt and Ostα (Figure 2).

**Figure 2.**
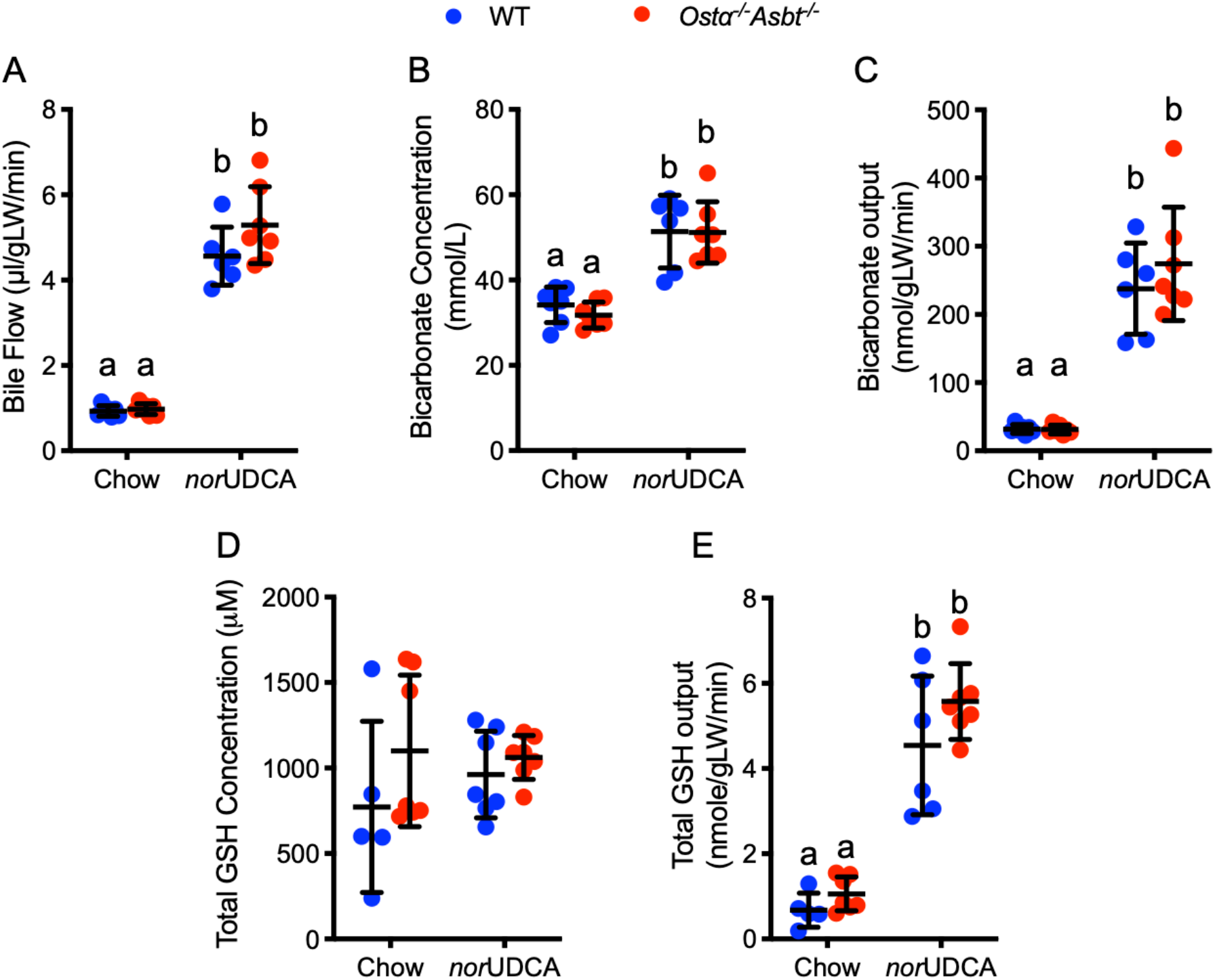
*nor*UDCA treatment increases bile flow and biliary bicarbonate and solute output in WT and *Ostα^−/−^Asbt^−/−^* mice. (**A**) Bile flow. (**B**) Biliary bicarbonate concentration. (**C**) Bicarbonate output. (**D**) Glutathione concentration. (**E**) Glutathione output. Mean + SD, n = 5-7 mice per group. Distinct lowercase letters indicate significant differences between groups (*P* < 0.05).

The negative findings for Asbt and Ostα-Asbt null mice do not exclude the potential involvement of other membrane transporters. We hypothesized that *nor*UDCA may act in a feed-forward fashion to induce hepatocyte or cholangiocyte expression of transporters involved in *nor*UDCA’s disposition, cholehepatic shunting, or mechanism of action. In order to identify potential candidates, RNA-Seq analysis was performed using livers from WT mice fed chow or *nor*UDCA-containing diets. Using a log2(fold-change) > 1 and multiple testing (FDR 5%), 1232 downregulated and 1087 upregulated genes were identified in *nor*UDCA-treated versus chow-fed WT mice. Narrowing our focus to membrane transporter gene expression revealed 80 Solute Carrier (*SLC*) family members, 16 ATP-binding cassette (*ABC*) family members, and 15 transporting P-type ATPases (*ATP*) family members (Figure 3A, 3B; Supplemental Figure 5A) that are differentially expressed in *nor*UDCA versus chow-fed control WT mice. Of these hepatic genes, expression of 30 *SLC*, 10 *ABC*, and 3 *ATP* transporters were significantly induced. Among the most highly induced transporter genes in *nor*UDCA-treated mice was *Slco1a4* (Oatp1a4; originally called Oatp2). Oatp1a4 is a sodium-independent facilitative uptake carrier with a broad substrate specificity that includes bile acids and has been reported to be expressed on the hepatocyte sinusoidal membrane (21).

**Figure 3.**
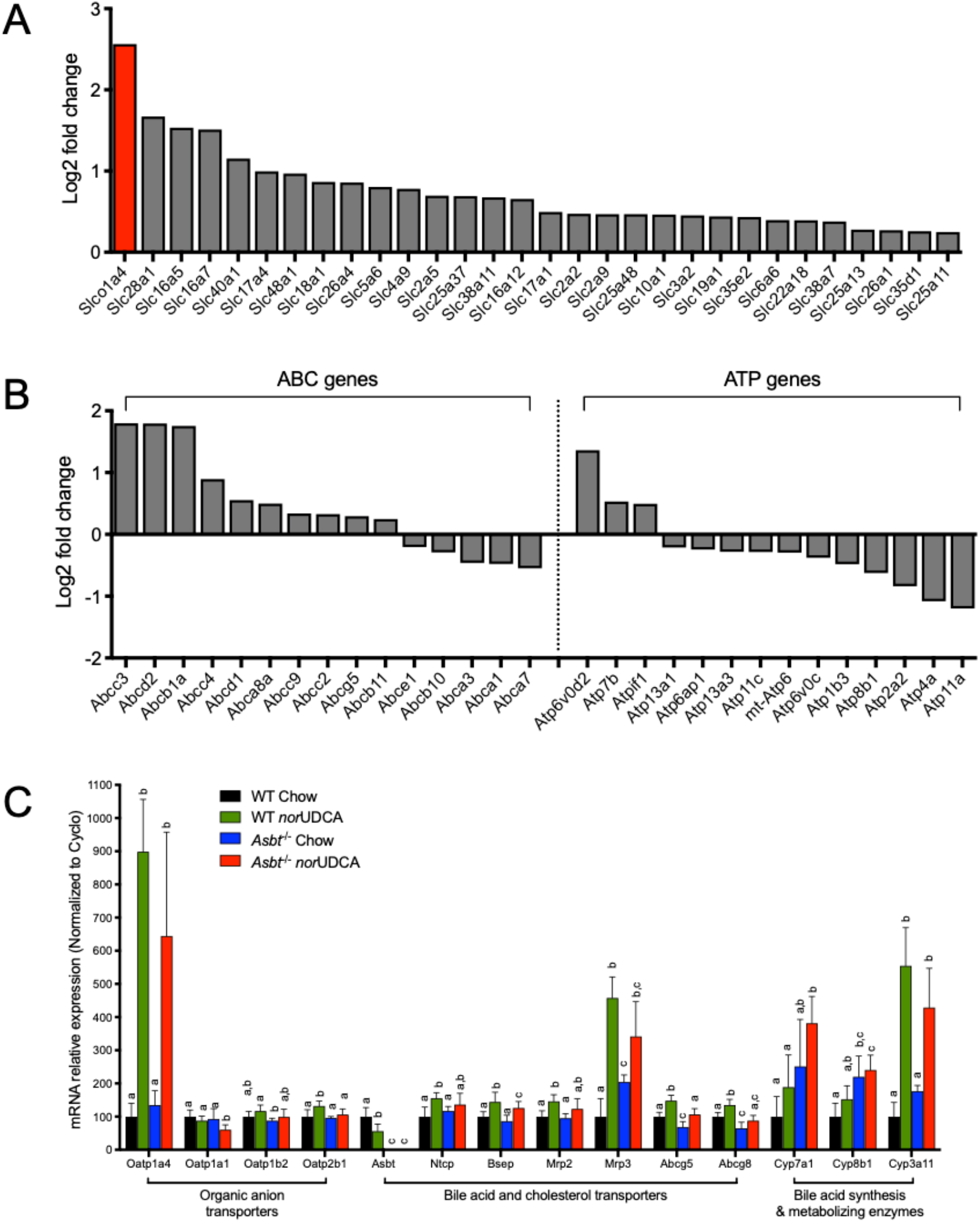
*nor*UDCA treatment alters expression of a limited number of hepatic transporter genes. RNA-Seq analysis of livers from WT mice fed chow or the *nor*UDCA-diet. (**A**) Differentially expressed *SLC* membrane transporter genes whose expression was significantly induced (*P* < 0.05; n = 6 per group) in *nor*UDCA-treated versus chow mice. (**B**) Differentially expressed *ABC* transporter and *ATP* P-type ATPase genes (*P* < 0.05; n = 6 per group) in the *nor*UDCA-treated versus chow mice. (**C**) Hepatic expression of the indicated transporters and bile acid-related biosynthesis or metabolizing enzymes in WT and *Asbt^−/−^* mice fed chow or the *nor*UDCA-containing diet for 7 days. RNA was isolated from livers of individual mice and used for real time PCR analysis. The mRNA expression was normalized using cyclophilin and the results for each gene are expressed relative to chow-fed WT mice (set at 100%). Mean + SD, n = 6-7 mice per group. Distinct lowercase letters indicate significant differences between groups (*P* < 0.05).

To further pursue the RNA-Seq findings, real-time PCR was used to measure mRNA expression of select transporters and genes critical for bile acid homeostasis (Figure 3C). Administration of *nor*UDCA induced Oatp1a4 mRNA expression by more than 6-fold in WT and *Asbt^−/−^* mice, whereas other hepatic OATP family genes, Oatp1a1, Oatp1b2 and Oatp2b1 were largely unaffected. With regard to other transporters involved in bile acid or cholesterol metabolism, hepatic expression of Asbt was decreased, whereas Ntcp, Bsep, Mrp2, Abcg5/8 were modestly increased and Mrp3 RNA levels were increased 3-4 fold (Figure 3C). In ileum, administration of *nor*UDCA tended to reduce mRNA expression of the bile acid-related genes Asbt, Fgf15, and Ostα-Ostβ, but had little effect on Ibabp expression (Supplemental Figure 5B). Notably, administration of norUDCA induced expression of a number of PXR-target genes, including *Slco1a4* (Oatp1a4), *Abcc3* (Mrp3), and *Cyp3a11*. These findings are in agreement with analysis of the RNA-Seq data, which identified pregnane X-receptor (PXR)-mediated direct regulation of xenobiotic metabolizing enzymes as one of the top-regulated pathways (Supplemental Figure 6A). To directly test the hypothesis that *nor*UDCA may be acting directly via PXR or other bile acid-activated nuclear receptors, the ability of *nor*UDCA to activate mouse PXR, human farnesoid X-receptor (FXR) and human vitamin D receptor (VDR) was examined in transfected human Huh7 cells. Over a range of concentrations up to 100 μM, *nor*UDCA failed to increase the activity of the PXR, FXR or VDR reporter plasmids (Supplemental Figure 6B).

The significant increase observed for hepatic Oatp1a4 expression raised the prospect that this transporter may be induced in a feed-forward fashion to facilitate hepatic clearance and cholehepatic shunting of *nor*UDCA. To directly test the hypothesis, the ability of *nor*UDCA to induce bile secretion and biliary bicarbonate output was examined in background (FVB) strain-matched WT and *Oatp1a/1b^−/−^* mice, in which *Slco1a1*, *Slco1a4*, *Slco1a5*, *Slco1a6* and *Slco1b2* have been excised by cre-mediated deletion of the *Slco1a/1b* gene cluster (22). The well-characterized *Oatp1a/1b^−/−^* mouse model was selected for these studies since the various Oatp1a and Oatp1b transporters display considerable overlap in their tissue expression and substrate specificity, and other members of the murine Oatp1a/1b subfamily could partially compensate for loss of Oatp1a4 alone. The experimental scheme and morphological response to *nor*UDCA feeding in *Oatp1a/1b^−/−^* mice are shown in Supplemental Figure 7. Administration of *nor*UDCA to the FVB background WT and *Oatp1a/1b^−/−^* mice for 7 days decreased body weight for both genotypes and increased the liver weight and liver to body weight ratio in the WT mice. Bile flow and biliary solute output are shown in Figure 4. Bile flow and biliary bicarbonate concentration, bicarbonate output, and pH were similar in the chow-fed WT and *Oatp1a/1b^−/−^* mice. Similar to WT C57BL/6J mice, administration of *nor*UDCA to WT FVB mice significantly increased bile flow by 4.7-fold, biliary bicarbonate concentration by 1.6-fold, and bicarbonate output by 7.5-fold as compared to chow-fed mice. In the *Oatp1a/1b^−/−^* mice, administration of *nor*UDCA induced a 5-fold increase in bile flow rate, 2-fold increase in biliary bicarbonate concentration, and 11-fold increase in bicarbonate output.

**Figure 4.**
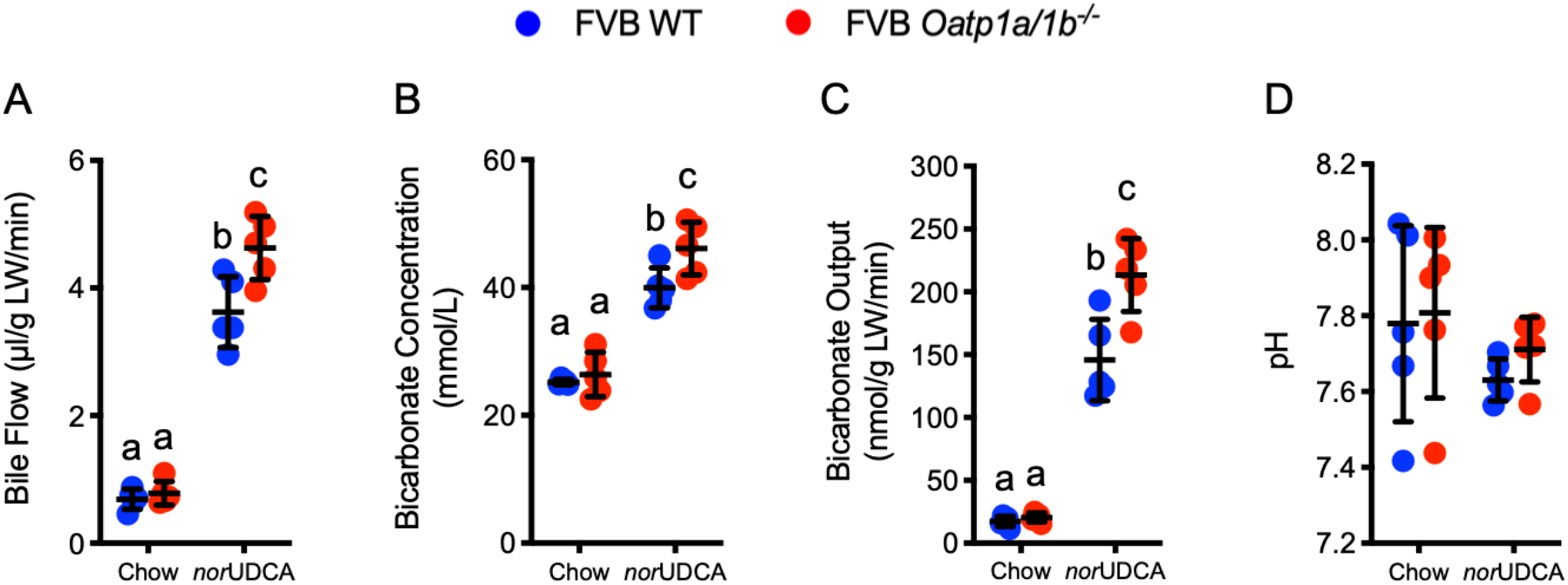
*nor*UDCA treatment increases bile flow and biliary bicarbonate and solute output in WT and *Oatp1a/1b^−/−^* mice. (**A**) Bile flow. (**B**) Biliary bicarbonate concentration. (**C**) Bicarbonate output. (**D**) Biliary pH. Mean + SD, n = 5 mice per group. Distinct lowercase letters indicate significant differences between groups (*P* < 0.05).

Administration of *nor*UDCA (2), ASBT inhibitors (23, 24), and bile acid sequestrants (25) have previously shown benefit in the *Abcb4/Mdr2^−/−^* mouse model of cholestasis and appear to involve both overlapping and complementary therapeutic mechanisms of action. Prompted by our observations of the *nor*UDCA-fed *Asbt^−/−^* mice, we next examined the effect of pharmacological interruption of the enterohepatic circulation of bile acids on the choleretic actions of *nor*UDCA by co-administering an ASBTi or bile acid sequestrant (Colesevelam). Colesevelam is a second generation bile acid sequestrant and non-absorbable polymer that binds bile acids through a combination of hydrophobic and ionic interactions and with a higher affinity than first generation sequestrants such as cholestyramine (26). Male WT mice were fed chow, chow supplemented with 0.006% (w/w) ASBTi, chow supplemented with 2% (w/w) colesevelam, or those diets plus 0.5% (w/w) *nor*UDCA for 7 days. The experimental scheme and morphological response to *nor*UDCA feeding is shown in Supplementary Figure 8. Administration of *nor*UDCA to mice for 7 days tended to reduce body weight, particularly when co-administered with an ASBTi (Supporting Fig. 8B, C), and increase the liver to body weight ratio (Supplementary Figure 8E). However, analysis of H&E-stained liver sections revealed no apparent histological differences between the treatment groups (Supplementary Figure 8F), and the plasma chemistries were not significantly different between the groups (Supplementary Table 3). The levels of bile acids excreted into the feces for each of the treatment group are shown in Supplementary Figure 8G. Administration of the ASBTi versus colesevelam resulted in a greater increase in the fecal bile acid content, however both treatments induced similar changes in fecal bile acid composition versus chow control mice (Supplementary Figure 8H). Administration of *nor*UDCA in the diet increased the fecal bile acid content in all treatment groups, due to increased excretion of the exogenous *nor*UDCA, as well as an increase in endogenous bile acids.

Similar to the findings for *Asbt^−/−^* mice, *nor*UDCA significantly increased bile flow by 3 to 4-fold, bicarbonate concentration by ~2-fold, and bicarbonate output by ~8-fold in WT mice when co-administered with an ASBTi (Figure 5A, 5B, 5C). Remarkably, *nor*UDCA also stimulated a similar bicarbonate-rich choleresis when co-administered with colesevelam. In agreement with the block in intestinal absorption of bile acids, biliary bile acid concentrations were reduced in ASBTi and colesevelam-treated versus control chow mice, however, administration of *nor*UDCA significantly increased biliary bile acid output in all the treatment groups (Figure 5F). The observation that co-administration of colesevelam did not attenuate the *nor*UDCA-induced hypercholeresis prompted an examination of the ability of colesevelam to bind *nor*UDCA versus endogenous bile acids in simulated small intestinal fluid. In accord with previous findings, colesevelam efficiently bound GCDCA and TDCA *in vitro* (27). However, under the same conditions, there was minimal binding of *nor*UDCA to colesevelam (Supplemental Figure 9) providing a potential explanation for the inefficacy of co-administered colesevelam to antagonize the actions of *nor*UDCA. In summary, pharmacological inhibition of intestinal bile acid absorption does not impede *nor*UDCA’s ability to induce a bicarbonate-rich hypercholeresis in mice.

**Figure 5.**
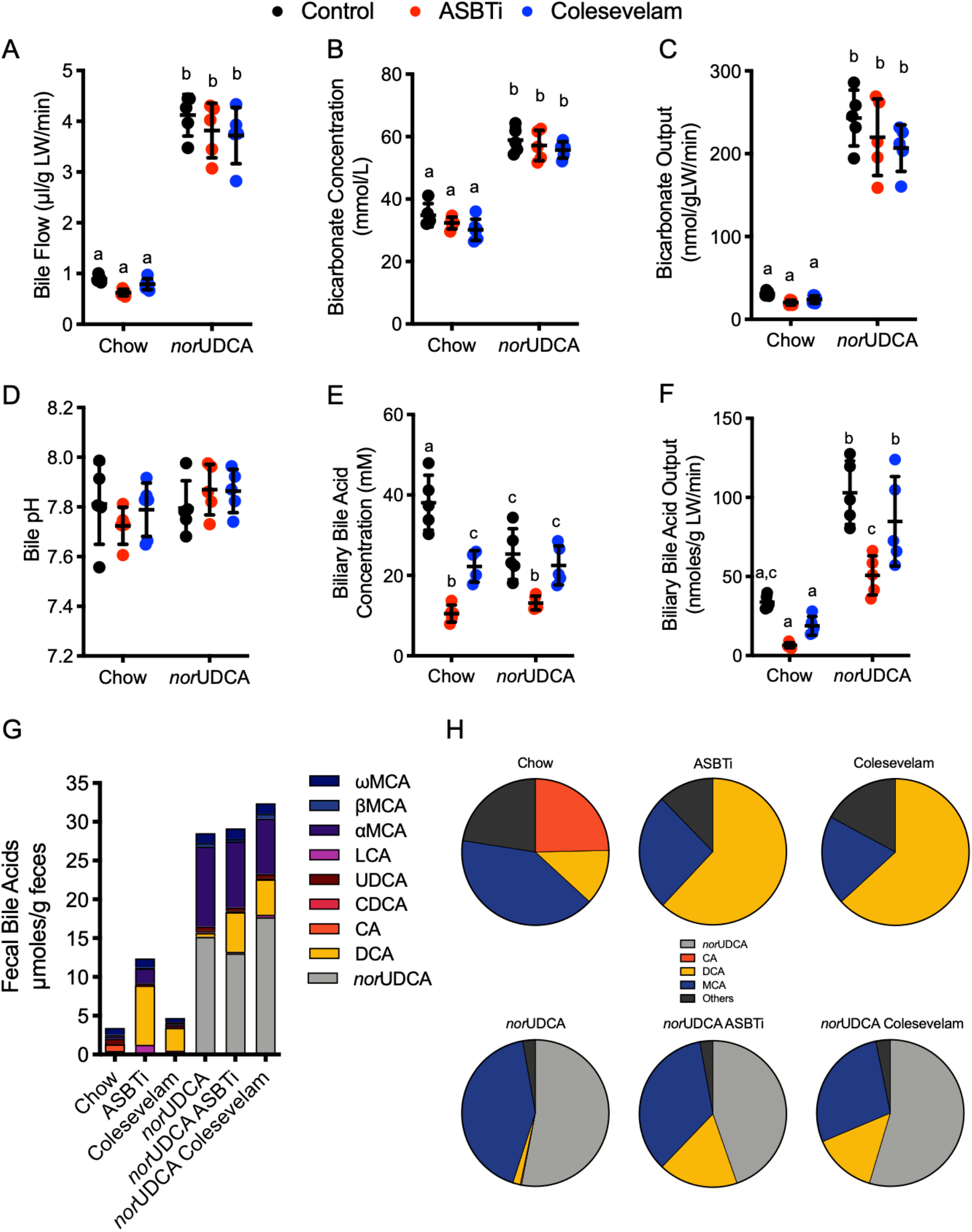
Pharmacological inhibition of intestinal bile acid absorption does not alter *nor*UDCA-induction of a bicarbonate-rich hypercholeresis in mice. (**A**) Bile flow. (**B**) Biliary bicarbonate concentration. (**C**) Bicarbonate output. (**D**) Biliary pH. (**E**) Biliary bile acid concentration. (**F**) Biliary bile acid output. (**G**) Fecal bile acids (mean values). (**H**) Fecal bile acid composition expressed as pie charts. Unless indicated, mean + SD are shown; n = 5 mice per group. Distinct lowercase letters indicate significant differences between groups (*P* < 0.05).

## Discussion

The major finding of this study is that the major bile acid transporters ASBT and OSTα-OSTβ are not required for orally-administered *nor*UDCA to stimulate a bicarbonate-rich hypercholeresis. The physiologic properties and metabolism of side chain-shortened C-23 *nor*-bile acids such as *nor*UDCA have been the subject of considerable study (1, 10, 11). Due to its higher critical micelle concentration (CMC) (17 mM) than many natural bile acids, *nor*UDCA is more likely present in monomeric rather than micellar form in bile (28). Based on their findings, Hofmann and coworkers proposed that C-23 *nor*-dihydroxy bile acids are sufficiently hydrophobic to be absorbed by passive diffusion (9). However, membrane transporters participate in the hepatic clearance of various hydrophobic drugs and endobiotics, including unconjugated C-24 bile acids (22, 29), and study of the contribution of individual bile acid and organic anion transporters to the transport of *nor*UDCA has been restricted to transfected cell-based models (16). The ASBT and OSTα-OSTβ play a central role in the intestinal reabsorption of bile acids and are expressed by the biliary epithelium (31–33). As such, it was possible that these major bile acid transporters could play a direct or indirect role in the absorption, cholehepatic shunting and actions of *nor*UDCA.

In agreement with previous studies, administration of *nor*UDCA significantly increased bile flow and bicarbonate concentration in WT mice, and similar results were observed for *Asbt^−/−^* and *Ostα^−/−^Asbt^−/−^* mice. Biliary bile acid output, typically lower in *Asbt^−/−^* versus WT mice, was significantly increased by administration of *nor*UDCA. This was driven primarily by the increase in bile flow, since the biliary total bile acid concentration was not significantly changed. However, *nor*UDCA administration significantly altered the biliary bile acid composition, with *nor*UDCA accounting for more than half the biliary bile acid species and a concomitant reduction in endogenous bile acid species. In *Asbt^−/−^* mice and mice treated with ASBT inhibitors, the biliary bile acid composition becomes more hydrophobic and enriched in TCA and TDCA. This is most likely due to increased hepatic TCA synthesis and an increased flux of TCA into the colon, where it is metabolized to deoxycholic acid (DCA), passively reabsorbed, and carried back to the liver for uptake, reconjugation and 7α-rehydroxylation of a portion of the TDCA to TCA (24, 30). Following administration of *nor*UDCA, the biliary endogenous bile acid composition in WT and *Asbt^−/−^* mice become remarkably similar. A potential mechanism for the decrease in TDCA is *nor*UDCA suppression of gut microbial cholic acid 7α-dehydroxylation since administration of *nor*UDCA also reduced the proportion of DCA in feces from *Asbt^−/−^* mice and WT mice treated with an ASBT inhibitor or bile acid sequestrant. Interestingly, there was an increase in fecal endogenous bile acids observed in WT mice treated with *nor*UDCA, which may be secondary to decreased ileal Asbt expression or weak inhibition of ileal Asbt activity. Using cell-based models, hydrophobic unconjugated bile acids, such as CDCA, DCA, UDCA, LCA, exhibited little apparent ASBT-mediated uptake over background, but are still able to compete for conjugated bile acid uptake (31).

We hypothesized that *nor*UDCA may induce hepatic expression of its own transporter(s) in a feed-forward fashion to facilitate cholehepatic shunting. RNA-Seq analysis revealed that *nor*UDCA induced expression of only a small subset of hepatic transporter genes, including several transporters involved in bile acid homeostasis, Oatp1a4, Mrp3, Mrp4, and Mdr1a. Interestingly, many of the genes induced by *nor*UDCA are PXR target genes, raising the prospect that *nor*UDCA may be acting directly or indirectly via PXR. However when interrogated, *nor*UDCA did not appear to directly activate mouse PXR or the bile acid sensing nuclear receptors, FXR or VDR as measured using nuclear receptor-luciferase reporter assays in transfected human hepatoma Huh7 cells. We focused our attention on Oatp1a4, since its hepatic expression was most highly induced in *nor*UDCA-fed WT mice. Oatp1a4 (originally called Oatp2) is expressed on the hepatocyte sinusoidal membrane, transports a variety of organic anions, and contributes to the hepatic clearance of steroid sulfates, bile acids, and drugs (32–34). However, the three most abundant hepatic Oatp isoforms in rodents, Oatp1a1, Oatp1b2 and Oatp1a4, exhibit overlapping substrate specificity (35), prompting us to use the *Oatp1a/1b^−/−^* mouse model lacking all 3 transporters for our studies (22). Similar to *Asbt^−/−^* and *Ostα^−/−^Asbt^−/−^* mice, loss of the Oatp1a/1b transporters did not impair the ability of norUDCA to stimulate a bicarbonate-rich hypercholeresis. This is in line with data obtained using cells stably expressing human liver OATPs, which failed to detect appreciable *nor*UDCA transport by human OATP1B1, OATP1B3 or OATP2B1 (16).

The finding that biliary bicarbonate concentration and bile flow increases in the *nor*UDCA-fed WT and various transporter knockout models is strongly consistent with the mechanism for cholehepatic shunting of *nor*UDCA proposed by Hofmann (1), however the study had a number of limitations. This included the use of a high pharmacological dose of *nor*UDCA administered in the diet. Although widely used for previous studies in mice (2–4), the higher dose may obscure the contribution of saturable carrier-mediated mechanisms to the absorption, disposition and actions of *nor*UDCA. Another limitation of the study is that only the unmodified *nor*UDCA was quantified and norUDCA metabolites such as the *nor*UDCA glucuronides were not measured (2, 10).

One of the more intriguing findings in this study is the ability of *nor*UDCA to increase bile flow and biliary bicarbonate when co-administered with an ASBTi or bile acid sequestrant. Based on our findings with the *Asbt^−/−^* mice, it was not surprising that the choleretic actions of *nor*UDCA was unaffected by an ASBTi. However, co-administration of *nor*UDCA with a bile acid sequestrant had not been previously examined. This contrasts with UDCA, whose interactions with bile acid sequestrants has been studied *in vitro*, in animal models, and in human subjects (36–38). In those studies, cholestyramine and colestimide efficiently bound and reduced the intestinal absorption of co-administered UDCA. As such, it was surprising that co-administration of colesevelam did not attenuate the choleretic actions of *nor*UDCA. Using *in vitro* assays, colesevelam bound *nor*UDCA poorly versus conjugated bile acids, however additional studies will be required to determine if this is a general property of all bile acid sequestrants. Although both ASBT inhibition and administration of bile acid sequestrants interrupt the enterohepatic circulation of bile acids, there are important differences regarding the mechanism of action by which they potentially improve features of cholestasis (23–25, 39). In the present study, ASBT inhibition reduces total biliary bile acid concentrations, yet in the presence of *nor*UDCA, bile flow was still induced. These findings indicate that pharmacological ileal ASBT inhibition not only doesn’t impair norUDCA’s positive effects on bile flow, but that in certain settings, may have beneficial synergistic effects in cholestatic models. This includes the reduced biliary bile acid concentration, more hydrophilic biliary bile acid composition, and elevated biliary bicarbonate concentration observed with *nor*UDCA plus ASBT inhibition versus ASBT inhibition alone.

Collectively, our findings suggest that *nor*UDCA does not require the major bile acid transporters, ASBT, OSTα-OSTβ or members of the OATP1a/1b family for absorption, cholehepatic shunting or to induce a bicarbonate-rich hypercholeresis. Importantly, these results also provide support for further investigation of the therapeutic potential of a combination of *nor*UDCA and blockers of the enterohepatic circulation of bile acids in cholestatic liver disease.

## Methods

### Materials

*nor*UDCA (24-nor-5β-cholan-23-oic acid) was received as a research gift from Dr. Falk Pharma to Dr. Michael Trauner. SC-435; (4R,5R)-5-[4-[4-(1-aza-4-azoniabicyclo[2.2.2]octan-4-yl)butoxy] phenyl]-3,3-dibutyl-7,8-dimethoxy-1,1-dioxo-4,5-dihydro-2H-1λ6C;6-benzothiepin-4-ol) was received as a research gift from Shire Pharmaceuticals. Colesevelam was provided by Dr. Alan Hofmann (University of California at San Diego).

### Animals

The *Asbt^−/−^* mice (*C57BL/6NJ-Slc10a2^tm1a(KOMP)Mbp^*; Asbt knockout-first, reporter-tagged insertion with conditional potential; Targeting Project CSD76540; https://www.komp.org/ProductSheet.php?cloneID=617849) were obtained from the Knockout Mouse Project (KOMP) Baylor College of Medicine Repository and colonies of *Asbt^−/−^* and matched WT mice are maintained at the Emory University School of Medicine. Characterization of the ileal and liver Asbt mRNA expression, fecal bile acid excretion, and bile acid pool size and composition in male and female WT and Asbt knockout-first mice is shown in Supplemental Figure 1. The matched background strain WT and *Ostα^−/−^Asbt^−/−^* mice were generated as described previously (20). Male Oatp1a/1b gene cluster knockout mice (*Oatp1a/1b^−/−^*) (FVB.129P2-Del(Slco1b2-Slco1a5)1Ahs) and background-matched WT FVB mice were purchased from Taconic Biosciences. For the ASBTi and colesevelam studies, WT male mice (C57BL/6J) mice were obtained from Jackson Labs.

### Animal treatments and bile flow measurements

All experiments were performed using male mice, 3 months of age (25-30 g body weight). The indicated genotypes were fed rodent chow (Envigo; Teklad custom Diet No. TD.160819; global 18% protein rodent diet) for 7 days. For the next 7 days, the mice were fed TD.160819 rodent chow or TD.160819 rodent chow containing the indicated combinations of 0.5% (w/w) *nor*UDCA, 0.006% ASBTi (SC-435; dose ~11 mg/kg/day), or 2% (w/w) colesevelam. The amount of *nor*UDCA, ASBTi, and colesevelam administered was selected based on published studies demonstrating sufficient doses to induce bile flow (*nor*UDCA) or disrupt the enterohepatic circulation of bile acids (ASBTi, colesevelam) (4, 25, 40). Based on an estimate of 3 g of diet consumed per day per 25 g body weight, the dose of *nor*UDCA was approximately 600 mg/kg/day. Bile flow was measured in mice as previously described (2). At the end of the bile collection period, blood was obtained by cardiac puncture to measure plasma chemistries. Portions of the liver were taken for histology and measurements of gene expression.

### Plasma biochemistries and biliary solute measurements

Plasma chemistries were measured at the Emory University Department of Animal Resources Quality Assurance and Diagnostic Laboratory. The bile samples were used immediately after isolation to measure HCO_3_-, pH, total CO_2_, Na, K, Cl, and glucose using a blood gas analyzer (i-STAT; Abbott Point of Care Inc) in the Clinical Pathology Laboratory, Emory University-Yerkes National Primate Research Center. Biliary glutathione concentrations were measured in the Emory University Pediatric Biomarkers Core Facility. Biliary bile acid, cholesterol and phospholipid concentrations were measured enzymatically as previously described (41, 42).

### Histological analysis

The liver segments were fixed in 10% neutral formalin (Sigma-Aldrich), embedded in paraffin and processed by Children’s Healthcare of Atlanta Pathology Services. Histological sections (5 μm) were cut and stained with H&E. The liver histology was assessed in a blinded fashion by a certified veterinary pathologist (S.G.).

### Bile acid measurements

To characterize the *Asbt^−/−^* mice, feces were collected from single-housed adult male and female mice over a 72-hour period. The total fecal bile acid content was measured by enzymatic assay (41, 43). Pool size was determined as the bile acid content of the small intestine, liver, and gallbladder removed from non-fasted mice (44, 45). Quantitative analysis of the biliary bile acids from chow or *nor*UDCA-fed mice was carried out at the Clinical Mass Spectrometry Laboratory at Cincinnati Children’s Hospital Medical Center as described (46). For the *nor*UDCA feeding studies, fecal samples were collected from the cages of group-housed mice with standard bedding at the end of the 7-day chow or *nor*UDCA feeding period. Fecal bile acid composition was determined using a Hewlett-Packard Agilent gas chromatography/mass spectrometer in the Pediatric Biomarkers Core Facility at Emory University as described (47).

### RNA-Seq analysis

Total RNA was extracted from frozen liver tissue using TRIzol reagent (Invitrogen, Carlsbad, CA). RNA-Seq libraries were prepared by Novogene Co., Ltd and sequenced on an Illumina HiSeq1000 system. Differential expression analysis was performed using the DESeq2 R package of Bioconductor (48). The resulting *P* values were adjusted using the Benjamini-Hochberg procedure to control for the false discovery rate (49). Differentially expressed genes with a fold change > 1.0 and adjusted *P* < 0.05 were selected for functional annotation (GEO series accession number: GSE145020). Pathway analysis of the RNA-Seq data was performed using MetaCore (GeneGo Inc, Saint Joseph, MI).

### Luciferase assays

Human liver cells (Huh7) were transfected with expression plasmids for a chimeric nuclear receptor encoding the ligand binding domain of mouse PXR fused to the DNA binding domain of GAL4 along with a 5x Upstream Activation Sequence (UAS)-luciferase reporter, or expression plasmids for human FXR or VDR along with FXR or VDR-responsive luciferase reporter plasmids. Ligand additions and measurements of luciferase activity were performed as described (40).

### In vitro Colesevelam bile acid binding assay

Bile acid binding to colesevelam was carried out as described (27).

### Statistical analyses

Mean values ± standard deviation are shown unless otherwise indicated. The data were evaluated for statistically significant differences using the Mann-Whitney test, the two-tailed Student’s t test, ANOVA and Tukey-Kramer honestly significant difference post-hoc test or Sidak’s multiple comparisons test (GraphPad Prism; Mountain View, CA). Differences were considered statistically significant at *P* < 0.05.

### Data availability

The liver RNA-Seq dataset is available from the GEO repository with the following accession number: GSE145020.

### Study approval

All animal experiments were approved by the Institutional Animal Care and Use Committees at Emory University.

## Author Contributions

J.L., J.K.T., S.J.K, C.D.F., M.T. and P.A.D designed the study and conceived the experiments. J.L, J.K.T., K.P., A.R., S.G, and P.A.D. performed experiments, collected results and analyzed the data. J.L., J.K.T. and P.A.D. managed the study. J.K.T., J.L., S.J.K, M.T. and P.A.D wrote the paper. J.L. and J.K.T. contributed equally to this work. J.L. initiated the project before J.K.T. joined the laboratory. After joining the lab, J.K.T. worked with J.L. to perform experiments, collect results and analyze data. JKT continued the project after J.L. left the laboratory to accept another position.

## Acknowledgements

We thank Shire Pharmaceutical for the research gift of the SC-435, Dr. Alan Hofmann (University of California, San Diego) for the research gift of the colesevelam, Drs. Kenneth Setchell and Wujuan Zhang (Cincinnati Children’s Clinical Mass Spectrometry Laboratory) for analysis of the mouse biliary bile acid samples, Dr. Steve Kliewer (University of Texas Southwestern Medical Center) for the PXR expression and reporter plasmids, and Dr. James Fleet (Purdue University) for the VDR expression plasmids. We also thank Ashley Bennet for assistance with the bile collections and Dr. Ivo P. van de Peppel for assistance with the nuclear receptor assays. We acknowledge the Emory Pediatrics Biomarkers Core for assistance with the bile acid and glutathione measurements, Children’s Healthcare of Atlanta Pathology Services for assistance processing tissues, and the Yerkes Nonhuman Primate Molecular Pathology Core for assistance with the histological analysis. Portions of this work were presented at the Annual Meeting of the American Association for the Study of Liver Disease in San Francisco, CA, 8-13 November 2018 and have appeared in abstract form, Hepatology 2018; 68: 46A. This research was supported by the National Institutes of Health, National Institute of Diabetes and Digestive and Kidney Diseases grants DK047987 (P.A.D.) and DK056239 (S.J.K.), the Meredith Brown Fund at Emory (S.J.K.), and Children’s Healthcare of Atlanta and Emory University’s Pediatric Biomarkers Core. JKT was supported by T32 GM008367. MT was supported by the Austrian Science Foundation (FWF) through projects F3517, F7310, and I2755.

