## Supplemental materials for "Active Enterohepatic Cycling is Not Required for the Choleretic Actions of 24-*nor*Ursodeoxycholic Acid in Mice"

**Supplemental Material**

**Abbreviations:** ABC, ATP-binding cassette; ALP, alkaline phosphatase; ALT, alanine aminotransferase; ASBT, apical sodium-dependent bile acid transporter; ASBTi, ASBT inhibitor; AST, aspartate aminotransferase; ATP, P-type ATPase; BSEP, bile salt export pump; CA, cholic acid; CHOL, cholesterol; DCA, deoxycholic acid; FGF15, fibroblast growth factor 15; FXR, farnesoid X-receptor; IBABP, ileal bile acid binding protein; LCA, lithocholic acid; Mdr2, multidrug resistance protein 2; OST, organic solute transporter; Ntcp, Na-taurocholate co-transporting polypeptide; OATP, organic anion transporting polypeptide; PSC, Primary Sclerosing Cholangitis; PXR, pregnane X-receptor; TBILI, total bilirubin; TCA, taurocholic acid; TCDCA, taurochenodeoxycholic acid; TDCA, taurodeoxycholic acid; THDCA, taurohyodeoxycholic acid; TαMCA, tauro-α-muricholic acid; TβMCA, tauro-β-muricholic acid; TωMCA, tauro-ω-muricholic acid; TUDCA, tauroursodeoxycholic acid; TP, total protein; VDR, vitamin D receptor; WT, wild type.

**Expanded Methods**

**Materials**

*nor*UDCA (24-*nor*-5β-cholan-23-oic acid) was received as a research gift from Dr. Falk Pharma to Dr. Michael Trauner. The purity of the *nor*UDCA was confirmed by HPLC. The ASBT inhibitor SC-435; (4R,5R)-5-[4-[4-(1-aza-4-azoniabicyclo[2.2.2]octan-4-yl)butoxy] phenyl]-3,3-dibutyl-7,8-dimethoxy-1,1-dioxo-4,5-dihydro-2H-1λ6-benzothiepin-4-ol) was received as a research gift from Shire Pharmaceuticals, and the ASBT inhibitory activity was confirmed using MDCK cells stably transfected with the human ASBT ($IC_{50}$ ~18 nM). Colesevelam was provided by Dr. Alan Hofmann (University of California at San Diego) and was received as a research gift from GelTex Pharmaceuticals/Genzyme.

**Animals and Tissue Collection**

All animal experiments were approved by the Institutional Animal Care and Use Committees at Emory University. The *Asbt*$^{-/-}$ mice (*C57BL/6NJ-Slc10a2*$^{tm1a(KOMP)Mbp}$; Asbt knockout-first, reporter-tagged insertion with conditional potential; Targeting Project CSD76540; https://www.komp.org/ProductSheet.php?cloneID=617849) were obtained from the Knockout Mouse Project (KOMP) Baylor College of Medicine Repository, and colonies of *Asbt*$^{-/-}$ and matched wild type (WT) mice generated from *Asbt*$^{+/-}$ crosses are maintained at the Emory University School of Medicine. The Asbt knockout-first mice were selected for these studies in place of the *Slc10a2*$^{tm1pda/J}$ whole body knockout mice (1) because they are in a pure C57BL/6NJ background and can be used to generate Asbt-floxed and tissue-specific null mice. Characterization of the ileal and liver Asbt mRNA expression, fecal bile acid excretion, and bile acid pool size in male and female WT and Asbt knockout-first mice is shown in **Supporting Fig. 1.** The *Ost*$\alpha^{-/-}$*Asbt*$^{-/-}$ and background-matched WT mice were generated as described previously (2). *Ost*$\alpha^{-/-}$*Asbt*$^{-/-}$ mice were selected for these studies in place of *Ost*$\alpha^{-/-}$ mice because inactivation of the Asbt protects *Ost*$\alpha^{-/-}$ mice from ileal injury and prevents/attenuates the associated adaptive changes including lengthening of the small intestine, ileal hypertophy, villous blunting, increased numbers of mucin-producing cells, and increased cell proliferation (2). These adaptive changes were predicted to complicate interpretation of the findings with regards to the role of Ost$\alpha$-Ost$\beta$ transport activity in the intestinal absorption and the choleretic actions of *nor*UDCA. Male Oatp1a/1b gene cluster knockout mice (*Oatp1a/1b*$^{-/-}$) (FVB.129P2-Del(Slco1b2-Slco1a5)1Ahs) and background-matched WT mice were purchased from Taconic Biosciences. For the ASBTi and colesevelam studies, WT male mice (C57BL/6J) were obtained from Jackson Labs. The mice were group-housed in ventilated cages (Super Mouse 750 Microisolator System; Lab Products) containing bedding (1/8" Bed-O-Cobbs; Andersons Lab Bedding Products) at temperature (22ºC) in the same 12h/12h light/dark cycle-controlled room of animal facility to minimize environmental differences and fed ad libitum rodent chow (13% of calories as fat; PicoLab Rodent Diet 20; LabDiet).

**Animal Treatment and Bile Flow Measurements**

All *nor*UDCA experiments were performed using male mice, 3 months of age (approximately 25-30 g body weight). The indicated genotypes were fed rodent chow (Envigo; Teklad custom Diet No. TD.160819; global 18% protein rodent diet) for 7 days. For the next 7 days, the mice were fed TD.160819 rodent chow or TD.160819 rodent chow containing the indicated combinations of 0.5% (w/w) *nor*UDCA, 0.006% ASBTi (SC-435; dose ~11 mg/kg/day), or 2% (w/w) colesevelam. The amount of *nor*UDCA, ASBTi, and colesevelam administered was selected based on published studies demonstrating these doses are sufficient to induce bile flow (*nor*UDCA) or disrupt the enterohepatic circulation of bile acids (ASBTi, colesevelam) (3-5). Based on an estimate of 3 g of diet consumed per day per 25 g body weight, the dose of *nor*UDCA administered is approximately 15 mg per 25 g mouse per day (600 mg/kg/day). At the end of the 7-day treatment period, the bile flow in the mice was measured as previously described (6). Briefly, non-fasted mice were anesthetized using an isoflurane/$O_2$ mixture, placed on a 37°C heating pad and maintained under isoflurane anesthesia during the procedure. Following laparotomy, the common bile duct was ligated, and gallbladder was cannulated. Residual gallbladder bile and hepatic bile secreted during the first 10 minutes of the collection period was discarded. Bile was then collected under a layer of mineral oil in pre-weighed tubes on ice for 120 minutes. Bile flow was determined gravimetrically and normalized to liver weight.

At the end of the bile collection period, blood was obtained by cardiac puncture to measure plasma chemistries. Liver wet weight was recorded at the time of necropsy. Portions of the liver were flash-frozen in liquid nitrogen and stored at -80º C for analysis of gene expression or fixed in 10% neutral formalin (Sigma-Aldrich) for embedding in paraffin and histological analysis. Small intestine was divided into 5 equal length segments and flushed with ice-cold phosphate-buffered saline (PBS) to remove luminal contents before weighing and flash-freezing in liquid nitrogen.

**Plasma Biochemistries and Biliary Solute Measurements**

Plasma chemistries, including total protein, albumin, alkaline phosphatase, alanine aminotransferase, aspartate aminotransferase, gamma-glutamyl transpeptidase, total bilirubin and cholesterol were measured at the Emory University Department of Animal Resources Quality Assurance and Diagnostic Laboratory. Immediately after isolation, the bile samples were used for determination of $HCO_3^-$, pH, total $CO_2$, Na, K, Cl, and glucose using a blood gas analyzer (i-STAT; Abbott Point of Care Inc) in the Clinical Pathology Laboratory, Emory University-Yerkes National Primate Research Center. Biliary glutathione concentrations were measured by HPLC in the Emory University Pediatrics Biomarkers Core Facility as described previously (7). Biliary bile acid and cholesterol concentrations were measured enzymatically as previously described (1, 8). In order to quantify biliary phospholipids, bile samples underwent chloroform-methanol extraction, followed by digestion with perchloric acid, and enzymatic assay to measure inorganic phosphate at the Wake Forest School of Medicine Lipid, Lipoprotein and Atherosclerosis Analysis Laboratory.

**Histological Analysis**

The liver tissue samples were fixed in 10% neutral formalin (Sigma-Aldrich) for 12 h and then stored in 70% (v/v) ethanol until processed for histology. Samples were embedded in paraffin and processed by Children’s Healthcare of Atlanta Pathology Services. Histologic sections (5 μm) were cut and stained with H&E. The liver histology was assessed in a blinded fashion by a certified veterinary pathologist (S.G.). Representative micrographs were collected using the Emory University Integrated Cellular Imaging Core.

**Bile Acid Measurements**

In order to characterize bile acid metabolism in the Asbt knockout-first mice, feces were collected from single-housed adult male and female mice over a 72-hour period. The total fecal bile acid content was measured by enzymatic assay (1, 9). Non-fasted mice were sacrificed and small intestine plus luminal contents, liver, and gallbladder were removed *en bloc* to measure the bile acid pool size. Total bile acids were extracted in ethanol (10), measured by enzymatic assay, and analyzed by HPLC as described (11). Individual bile acid species were identified using an evaporative light scatter detector (Alltech ELSD 800). Bile acids were quantified by comparison to known amounts of authentic standards purchased from Steraloids (Newport, RI). Quantitative analysis of the biliary bile acids from chow or *nor*UDCA-fed mice was carried out at the Clinical Mass Spectrometry Laboratory at Cincinnati Children's Hospital Medical Center. Briefly, biliary bile acids were extracted on a reverse-phase/solid-phase cartridge and analyzed using a Waters Quattro Premier XE triple quadruple mass spectrometer interfaced with Aquity UPLC system. Quantification of bile acids was based on a validated isotope dilution mass spectrometry method (12). Calibration curves were built with 20 of the mouse bile acids using the single ion recording (SIR) function on the mass spectrometer (LLOQ = 5 ng/ml; ULOQ = 1000 ng/ml). The bile acid Hydrophobic index was calculated according to Heuman using a value of -0.68 for the *nor*UDCA anion (13, 14).

For the *nor*UDCA feeding studies, fecal samples were collected from cages of group-housed mice with standard bedding at the end of the 7-day chow or *nor*UDCA feeding period. The fecal bile acid composition was determined using a Hewlett-Packard Agilent gas chromatography-mass spectrometer (GC/MS) in the Department of Pediatrics Biomarkers Core Facility at Emory University. Briefly, 50 mg of crushed dried feces mixed with 10 nmol of internal standard, 5β-Cholanic acid 7α, 12α-diol (Steraloids), resuspended in 75% methanol plus 250 mM NaOH and heated at 80°C for 2 h. After centrifugation, the bile acids were extracted on a reverse-phase/solid-phase cartridge (C-18 columns; Supelco) and dried down under nitrogen. The bile acids were then methylated by resuspension in acetyl-chloride (Sigma-Aldrich)/methanol (1:20; v:v) and heating at 55ºC for 20 min. For derivatization, the samples were dried under nitrogen, resuspended in N,O-Bis(trimethylsilyl)trifluoroacetamide (Sigma-Aldrich)/ pyridine (Sigma-Aldrich)/ chlorotrimethyl-silane (Sigma-Aldrich) (5:5:0.1; v:v:v), and incubated for 1 h at room temperature. The derivatized bile acids were then dried at room temperature and resuspended in heptane for GC/MS analysis and compared to authentic standards for *nor*UDCA, UDCA, CDCA, LCA, DCA, CA, αMCA, βMCA, and ωMCA (15).

**RNA-Seq Analysis**

Total RNA was extracted from frozen liver tissue using TRIzol reagent (Invitrogen, Carlsbad, CA). RNA-Seq libraries were prepared by Novogene Co., Ltd and sequenced on an Illumina HiSeq1000 system. The original data obtained from the high throughput sequencing platforms were transformed to sequenced reads by base calling. Raw data are recorded in a FASTQ file which contains sequenced reads and corresponding sequencing quality information. Sequences were aligned to the mouse genome using STAR (16). The number of reads that mapped to each gene was used to quantify RNA expression, which is indicated as fragments per kilobase or transcript sequence per million base pairs sequenced (17). Differential expression analysis was performed using the DESeq2 R package of Bioconductor (18). The resulting $P$ values were adjusted using the Benjamini-Hochberg procedure to control for the false discovery rate (19). Differentially expressed genes with a fold change > 1.0 and adjusted $P < 0.05$ were selected for functional annotation (GEO series accession number: GSE145020). Pathway analysis of the RNA-Seq data was performed using MetaCore (GeneGo Inc, Saint Joseph, MN) with a threshold fold change of 2 and $P$-value of 0.05 to indicate a statistically significant difference.

**Luciferase Assay**

Human liver cells (Huh7) were seeded in a 24-well plate at 1 X $10^5$ cells/well in monolayers and cultured at 37°C and 5% $CO_2$ in modified Eagle's minimal essential medium (EMEM) supplemented with 10% fetal bovine serum (FBS) and 1% penicillin-streptomycin. The following day, cells at ~85% confluency were transfected using Lipofectamine3000 (Invitrogen) to introduce a control renilla expression plasmid (0.005 µg), where the renilla luciferase gene is driven by a constitutively active promoter from the HSV thymidine kinase gene to monitor transfection efficiency, along with the following plasmid combinations: 1) a chimeric nuclear receptor encoding the ligand binding domain of mouse PXR fused to the DNA binding domain of GAL4 (pGAL4-mPXR-LBD, 0.25 µg) and a 5x Upstream Activation Sequence (UAS)-luciferase reporter (pFR-Luc, 0.025 µg), 2) expression plasmids for human FXR (0.025 µg), RXRα (0.025 µg), and an FXR-responsive luciferase reporter (pECRE-Luc, 0.25 µg), or 3) expression plasmids for human VDR (0.025 µg), RXRα (0.025 µg), and a VDR-responsive luciferase reporter (hCYP24-luc, 0.25 µg). The next day, triplicate wells were incubated in charcoal stripped FBS-containing medium plus vehicle (DMSO), different concentrations of positive control (pregnenolone 16α-carbonitrile, a mouse PXR ligand; GW4064, a synthetic FXR ligand, or 25-hydroxy-vitamin D, a vitamin D receptor ligand), or different concentrations of *nor*UDCA. After a 24 hour treatment, cells were washed in 1X PBS, harvested and processed using Luciferase assay reagents (Dual-Luciferase Reporter Assay System) from Promega. The bioluminescence was monitored using a luminometer (Synergy HTX, Biotek). The results were normalized to TK-renilla activity and presented as relative fold change over the DMSO vehicle group.

### *In vitro* Colesevelam Bile Acid Binding Assay

Bile acid binding to colesevelam was carried out as described (20). Briefly, bile acids were dissolved in simulated intestinal fluid (50 mM $KH_2PO_4$, 22.4 mM NaOH, pH 6.8) at a concentration of 2 mM for TDCA or 6 mM for GCDA and norUDCA, and pre-incubated for 30 min on a rotator at 37°C prior to addition to 7.5 mg of colesevelam hydrochloride. Samples were withdrawn at 0, 0.5, 1, 2, 3, 4 and 5 h, filtered through a 0.2 μm syringe filter, and the filtrate was analyzed by enzymatic assay to determine the free bile acid concentration. The difference between the initial (0 h) and free bile acid concentrations was used to calculate the amount of the bile acid bound per mg of colesevelam hydrochloride.

**Statistical Analyses**

Mean values ± standard deviation are shown unless otherwise indicated. The data were evaluated for statistically significant differences using the Mann-Whitney test, the two-tailed Student's t test, ANOVA (with a Tukey-Kramer honestly significant difference post-hoc test) or Sidak's multiple comparisons test (GraphPad Prism; Mountain View, CA). Differences were considered statistically significant at $P < 0.05$.

**Table 1. Physical and Permeability Properties of UDCA versus *nor*UDCA**

| | UDCA | *nor*UDCA |
|---|---|---|
| **Formula** | C24H40O4[1] | C23H38O4[1] |
| **Molecular Weight** | 392.56[1] | 378.553[1] |
| **xLogP3** | 4.9[1] | 4.6[1] |
| **Hydrogen Bond Donor** | 3[1] | 3[1] |
| **Hydrogen Bond Acceptor** | 4[1] | 4[1] |
| **Critical Micelle Concentration (mM)** | 7[2] | 17[3] |
| **Apparent Permeability (x $10^4$) (cm/sec in perfused rat jejunum)** | 0.67[4] | 0.64[4] |

1. Values computed by PubChem XLogP3 3.0, Cactvs 3.4.6.11.
2. Hofmann A and Roda A, Hepatology 1984: 25: 1477-1489.
3. Roda A, Hofmann AF, Mysels KJ. J Biol Chem 1983; 258: 6362-6370.
4. Dupas J-L et al, Lipids 2018; 53: 465-468.

### Table 2. Plasma Chemistries in Chow and *nor*UDCA Diet-fed Wild Type and *Asbt*$^{-/-}$ Mice

| Variable | WT (n) | | *Asbt*$^{-/-}$ (n) | |
|---|---|---|---|---|
| | Chow (6) | *nor*UDCA (7) | Chow (7) | *nor*UDCA (7) |
| **Albumin (g/dL)** | 1.8 ± 0.8$^{a}$ | 3.1 ± 0.5$^{a,b}$ | 2.6 ± 0.5$^{b}$ | 2.9 ± 0.4$^{b}$ |
| **Alkaline phosphatase (U/L)** | 38 ± 12 | 37 ± 14 | 33 ± 12 | 45 ± 5 |
| **Alanine aminotransferase (U/L)** | 85 ± 59 | 61 ± 27 | 120 ± 49 | 95 ± 39 |
| **Aspartate aminotransferase (U/L)** | 116 ± 67$^{a,b}$ | 99 ± 28$^{a}$ | 199 ± 70$^{b}$ | 146 ± 48$^{a,b}$ |
| **Cholesterol (mg/dL)** | 95 ± 27$^{a}$ | 151 ± 30$^{a,b}$ | 109 ± 18$^{b}$ | 148 ± 41$^{b}$ |
| **Gamma-glutamyl transpeptidase (U/L)** | 9.6 ± 4.6$^{a}$ | 15.4 ± 4.9$^{a,b}$ | 15.4 ± 2.8$^{a,b}$ | 17.1 ± 3.8$^{b}$ |
| **Total bilirubin (mg/dL)** | 3.0 ± 1.3$^{a,b}$ | 2.6 ± 0.9$^{a}$ | 3.6 ± 0.6$^{b}$ | 2.8 ± 0.7$^{a,b}$ |
| **Total protein (g/dL)** | 3.6 ± 0.6 | 4.6 ± 0.9 | 3.6 ± 0.9 | 4.6 ± 0.5 |

Values are expressed as the mean ± SD. The number of mice per group are indicated (n). Values with different superscript letters are significantly different ($P < 0.05$) by ordinary two-way ANOVA and Sidak’s multiple comparisons test.

**Table 3. Plasma Chemistries in Wild Type Mice after co-administration of *nor*UDCA with an ASBTi or Colesevelam**

| Variable | Control | ASBTi | Colesevelam | *nor*UDCA | *nor*UDCA ASBTi | *nor*UDCA Colesevelam |
|---|---|---|---|---|---|---|
| **Albumin (g/dL)** | 3.3 ± 0.2 | 3.0 ± 0.2 | 3.1 ± 0.2 | 3.4 ± 0.2 | 3.0 ± 0.7 | 3.5 ± 0.2 |
| **ALP (U/L)** | 6.4 ± 6.1 | 1.6 ± 2.2 | 8.8 ± 7.2 | 6.4 ± 5.4 | 4.0 ± 2.8 | 6.4 ± 4.6 |
| **ALT (U/L)** | 96 ± 27 | 96 ± 27 | 118 ± 46 | 74 ± 23 | 143 ± 96 | 97 ± 53 |
| **AST (U/L)** | 176 ± 60 | 170 ± 46 | 247 ± 145 | 90 ± 25 | 193 ± 141 | 115 ± 33 |
| **CHOL (mg/dL)** | 123 ± 13$^{a}$ | 125 ± 10$^{a}$ | 110 ± 9$^{a}$ | 171 ± 14$^{b}$ | 130 ± 44$^{a,b}$ | 148 ± 10$^{a,b}$ |
| **GGT (U/L)** | 19.2 ± 4.4 | 16.8 ± 3.3 | 14.4 ± 3.6 | 18.4 ± 6.1 | 15.2 ± 3.3 | 16.8 ± 3.3 |
| **TBILI (mg/dL)** | 2.6 ± 0.3$^{a}$ | 3.4 ± 0.4$^{a}$ | 2.6 ± 0.5$^{a}$ | 2.0 ± 0.4$^{b}$ | 3.1 ± 1.0$^{a}$ | 1.9 ± 0.4$^{b}$ |
| **TP (g/dL)** | 4.8 ± 0.3 | 4.3 ± 0.3 | 4.7 ± 0.4 | 5.1 ± 0.4 | 4.4 ± 1.2 | 5.4 ± 0.4 |

Values are expressed as the mean ± SD (n = 5 mice per group). Values with different superscript letters are significantly different ($P < 0.05$) by ordinary two-way ANOVA and Sidak's multiple comparisons test. ALP, alkaline phosphatase; ALT, alanine aminotransferase; AST, aspartate aminotransferase; CHOL, cholesterol; GGT, gamma-glutamyl transpeptidase; TBILI, total bilirubin; TP, total protein.

**Supplemental Figure Legends**

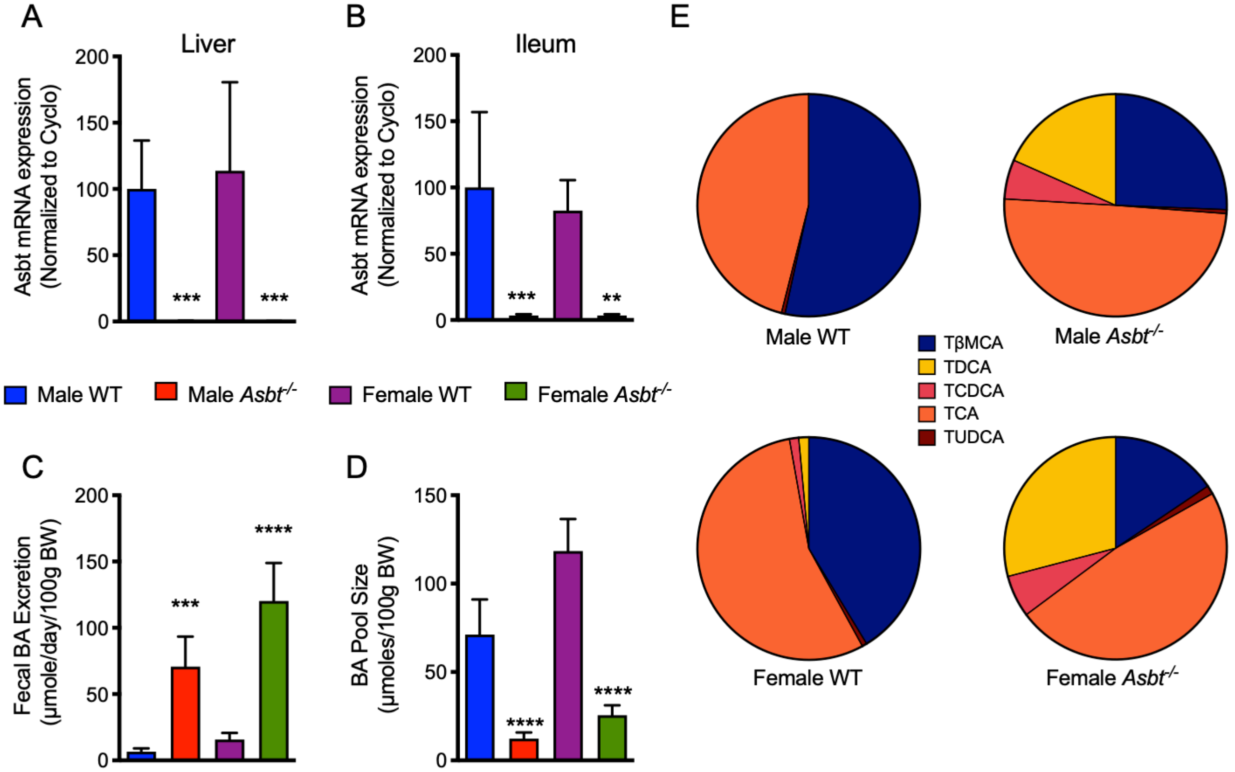


**Supplemental Figure 1. Asbt expression, bile acid excretion, bile acid pool size, and pool composition in wild type and Asbt knockout-first (*Asbt*$^{-/-}$) mice.** (**A**) Asbt mRNA levels in liver and (**B**) ileum of male and female WT and *Asbt*$^{-/-}$ mice. RNA was isolated from individual mice and used for real-time PCR analysis. The mRNA expression was normalized to cyclophilin and are shown relative to male WT mice (set at 100%) for each tissue. (**C**) Fecal bile acid excretion in male and female WT and *Asbt*$^{-/-}$ mice. The data are expressed as µmoles/per day per 100 g body weight. (**D**) Bile acid pool size in in male and female WT and *Asbt*$^{-/-}$ mice. The data are expressed as µmoles per 100 g body weight. (**E**) Bile acid pool composition in male and female WT and *Asbt*$^{-/-}$ mice. All mice were 3 months of age. Mean values $\pm$ SD are shown (n = 5 mice per group). Asterisks indicate significant differences (**$P < 0.01$, ***$P < 0.001$, ****$P < 0.0001$) between genotypes for each sex.

Asbt mRNA expression is reduced to background levels in liver and ileum of *Asbt*$^{-/-}$ mice. Fecal bile acid excretion was increased by approximately 8 to 10-fold in male and female *Asbt*$^{-/-}$ versus WT mice. The increased fecal bile acid excretion was associated with approximately 80 percent reductions in the bile acid pool size of male and female *Asbt*$^{-/-}$ mice. Analysis of the bile acid pool composition by HPLC revealed a large reduction in the proportion of 6-hydroxylated bile acid species and concomitant increase in the proportion of taurocholic acid and its gut microbiota-derived product taurodeoxycholic acid. These changes in fecal bile acid excretion, bile acid pool size and composition in the C57BL/6NJ Asbt knockout-first mice recapitulates the major bile acid phenotype of the previously characterized Asbt knockout mouse model (1). BA, bile acid; BW, body weight; TβMCA, tauro-beta-muricholic acid; TCA, taurocholic acid; TCDCA, taurochenodeoxycholic acid; TDCA, taurodeoxycholic acid; TUDCA, tauroursodeoxycholic acid; WT, wild type.

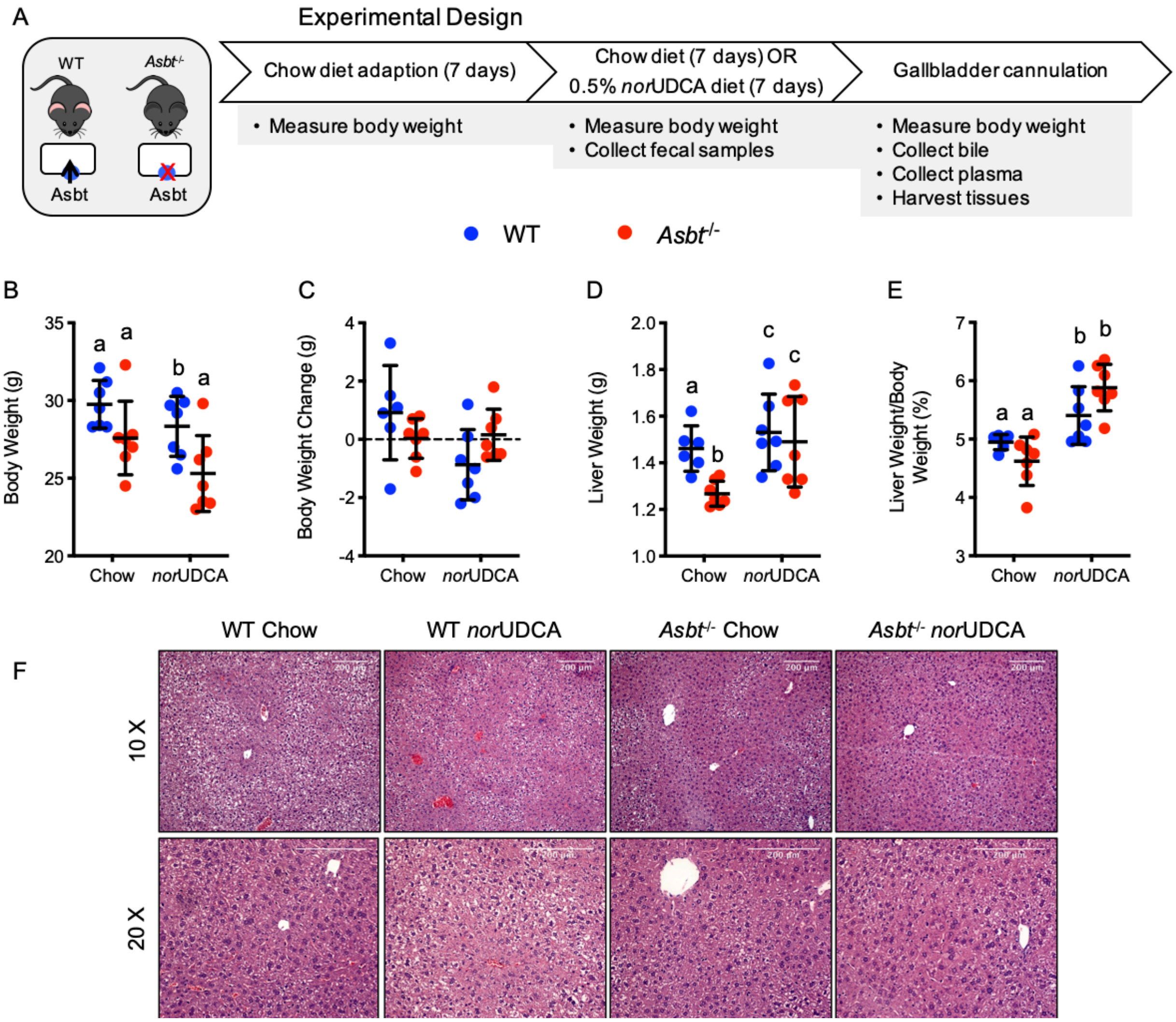


**Supplemental Figure 2. Experimental scheme and morphological response to *nor*UDCA treatment in WT and *Asbt*$^{-/-}$ mice.** (**A**) Experimental design. (**B**) Body weight after 7-days of feeding chow or chow plus 0.5% *nor*UDCA. (**C**) Body weight change after 7-days of feeding chow or chow plus 0.5% *nor*UDCA. (**D**) Liver weight. (**E**) Liver to body weight ratio. Mean $\pm$ SD, n = 6-7 mice per group. Distinct lowercase letters indicate significant differences between groups ($P < 0.05$). (**F**) Hematoxylin and eosin-stained liver sections (original magnification 10x or 20x) from the indicated genotypes and treatments groups. *Scale bar*, 200 µm.

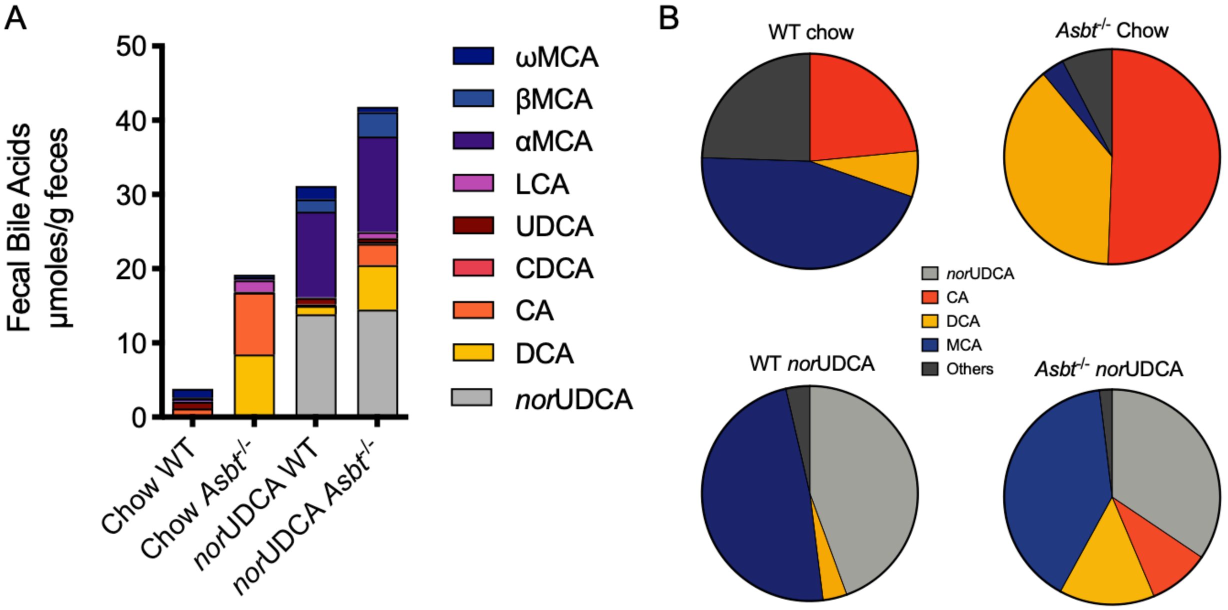


**Supplemental Figure 3. Effect of *nor*UDCA treatment on the fecal bile acid profile in WT and *Asbt*$^{-/-}$ mice.** (**A**) *nor*UDCA treatment increased the excretion of endogenous bile acids and the amount of 6-hydroxylated bile acids in feces. (**B**) Pie charts for the fecal bile acid profiles. Mean values are shown for fecal samples collected from cages of group-housed mice (n = 6-7 mice per genotype and treatment condition).

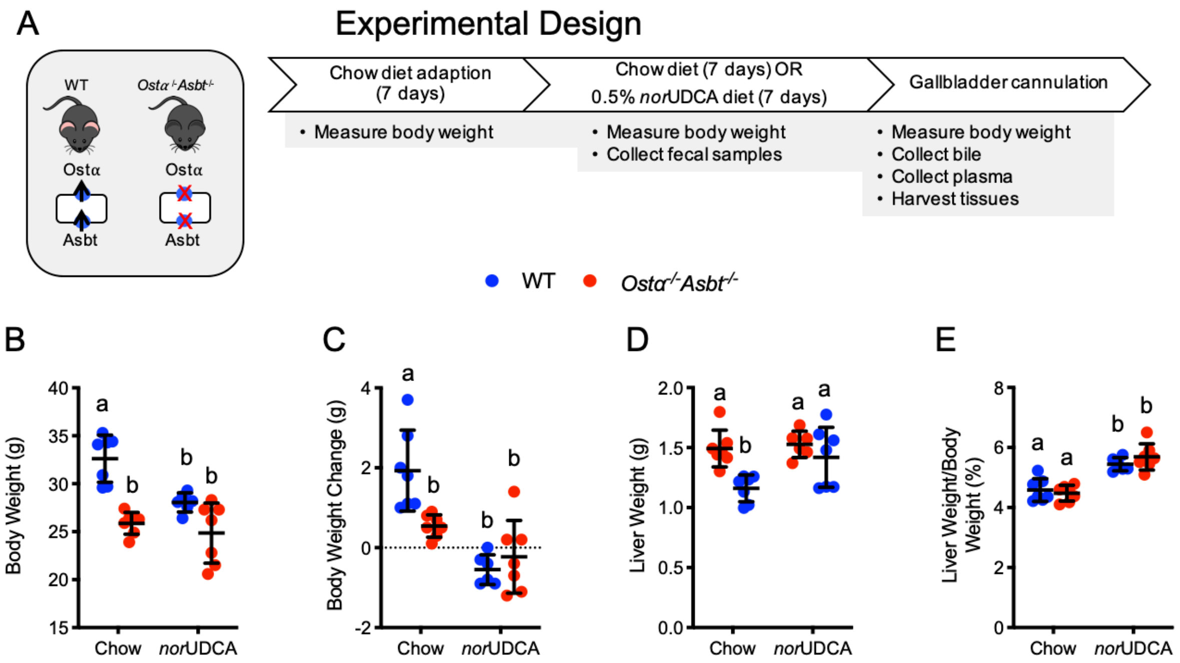


**Supplemental Figure 4. Experimental scheme and morphological response to *nor*UDCA treatment in WT and *Ostα$^{-/-}$Asbt$^{-/-}$* mice.** (**A**) Experimental design. (**B**) Body weight after 7-days of feeding chow or chow plus 0.5% *nor*UDCA. (**C**) Body weight change after 7-days of feeding chow or chow plus 0.5% *nor*UDCA. (**D**) Liver weight. (**E**) Liver to body weight ratio. Mean $\pm$ SD, n = 5-7 mice per group. Distinct lowercase letters indicate significant differences between groups ($P$ < 0.05).

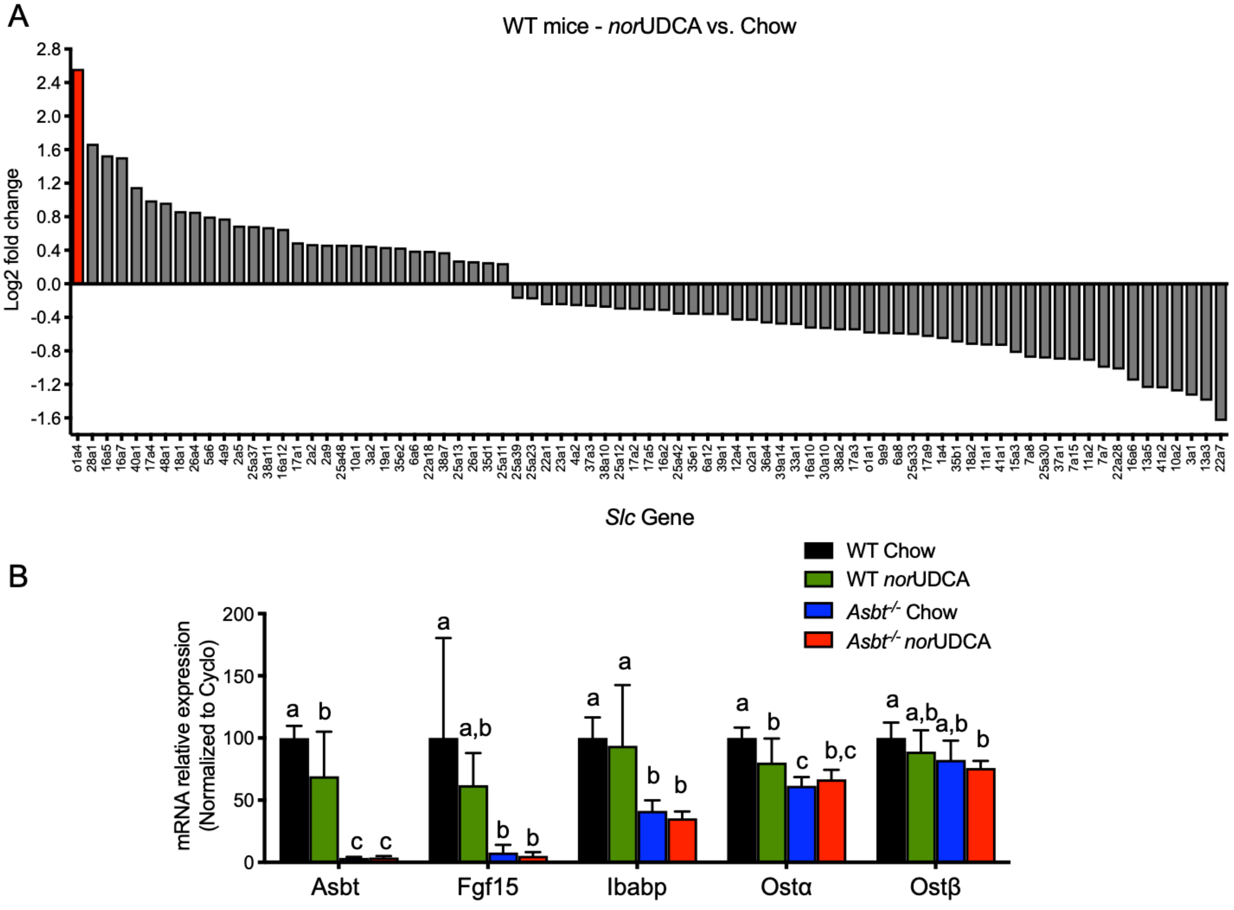


**Supplemental Figure 5. Effect of *nor*UDCA treatment on hepatic and ileal gene expression.** (**A**) RNA-Seq analysis of livers from WT mice fed chow or the *nor*UDCA-containing diet. Differentially expressed *SLC* membrane transporter genes ($P$ < 0.05; n = 6 per group) in the *nor*UDCA-treated versus chow mice are shown. (**B**) Ileal expression of the indicated bile acid-related genes in WT and *Asbt*$^{-/-}$ mice fed chow or the *nor*UDCA-containing diet for 7 days. RNA was isolated from ileum of individual mice and used for real-time PCR analysis. The mRNA expression was normalized using cyclophilin and the results for each gene are expressed relative to chow-fed WT mice for each gene. Mean ± SD, n = 6-7 mice per group. Distinct lowercase letters indicate significant differences between groups ($P$ < 0.05).

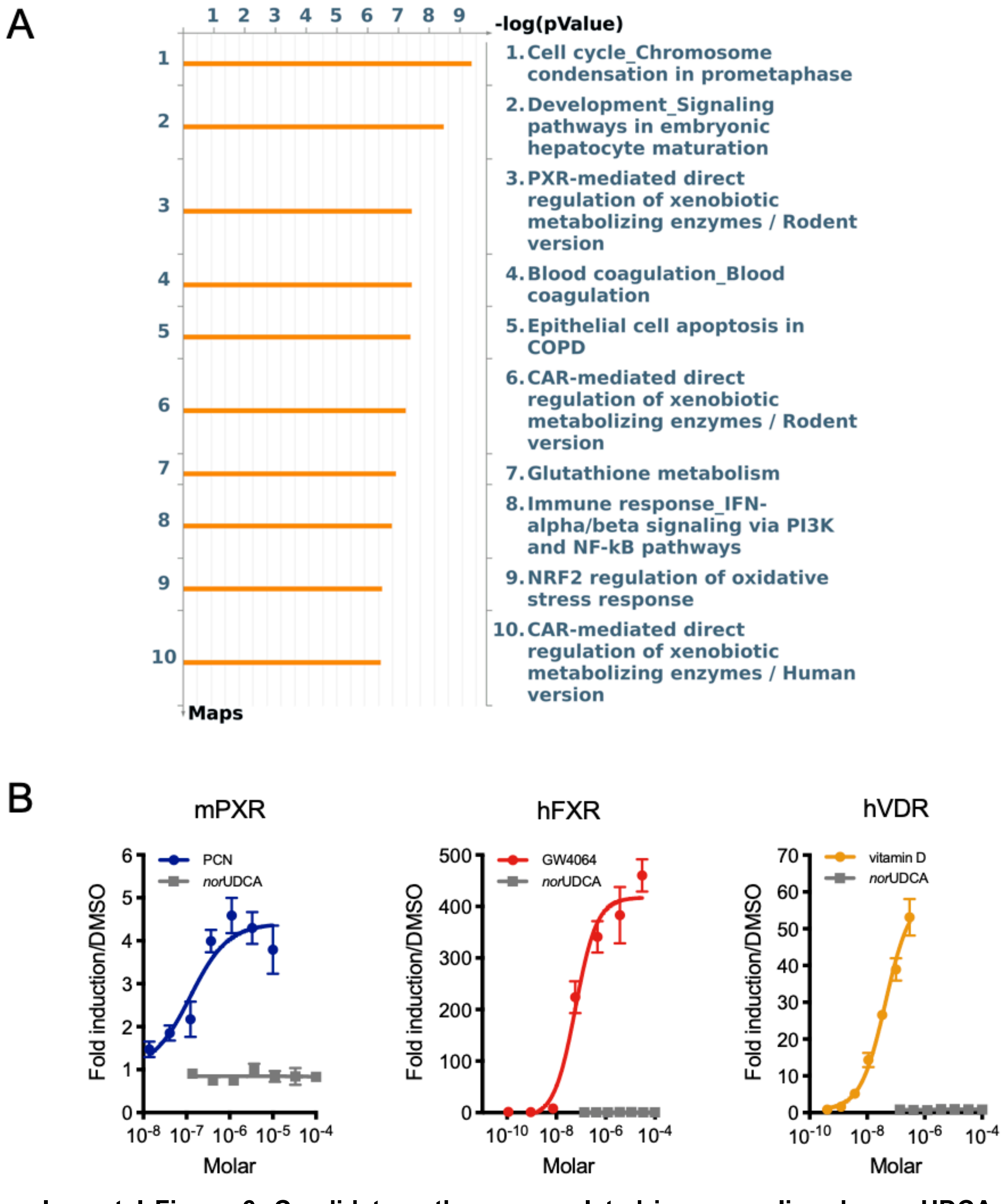


**Supplemental Figure 6. Candidate pathways regulated in mouse liver by *nor*UDCA.** (**A**) Pathway analysis of liver gene expression changes in *nor*UDCA treated versus chow-fed mice using a threshold fold change of 2 and *P* value of 0.05. The top 10 pathways and -log(pValue) are shown. (**B**) *nor*UDCA does not directly activate mouse PXR, human FXR or human VDR in transfected Huh7 cells. Huh7 cells were transfected with the indicated nuclear receptor expression and reporter plasmid combinations. One day following transfection, the cells were treated with the indicated concentrations of positive control ligand or *nor*UDCA for 24 h prior to luciferase activity measurement. The results were normalized to TK-renilla activity and presented as relative fold change versus vehicle (DMSO). Mean values ± SD are shown.

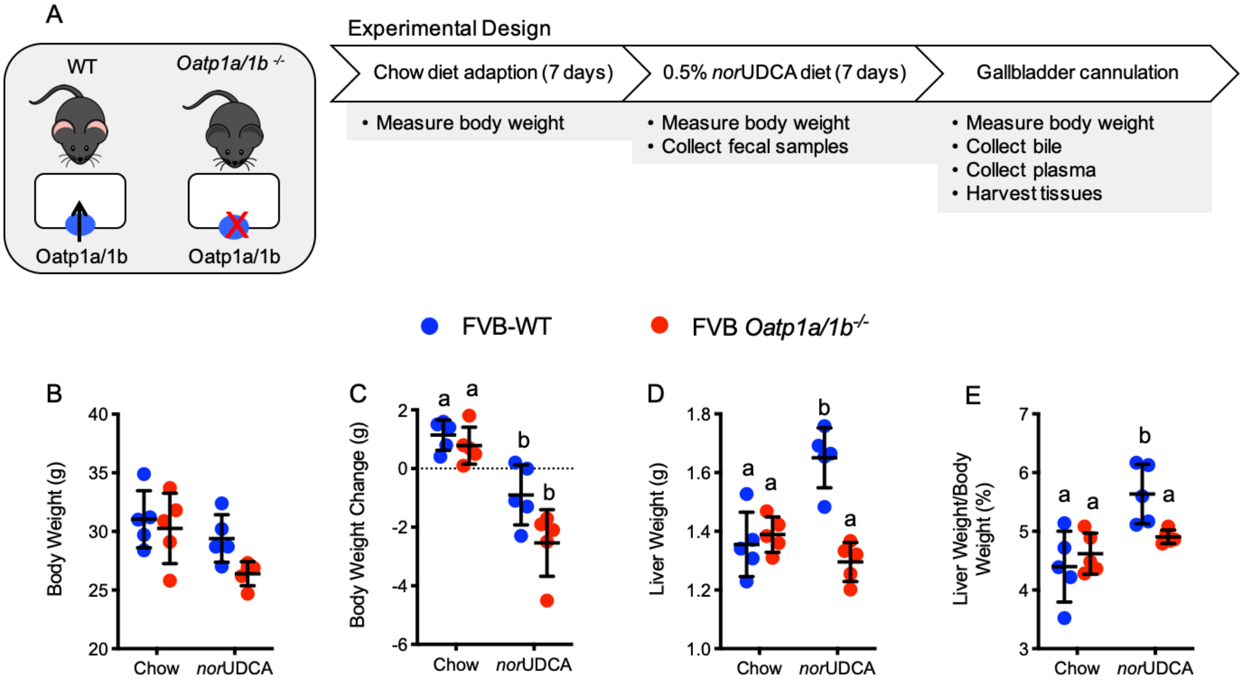


**Supplemental Figure 7. Experimental scheme and morphological response to *nor*UDCA treatment in WT and *Oatp1a/1b*$^{-/-}$ mice.** (**A**) Experimental design. (**B**) Body weight. (**C**) Body weight change. (**D**) Liver weight. (**E**) Liver to body weight ratio. Mean ± SD, n = 5 mice per group. Distinct lowercase letters indicate significant differences between groups ($P < 0.05$).

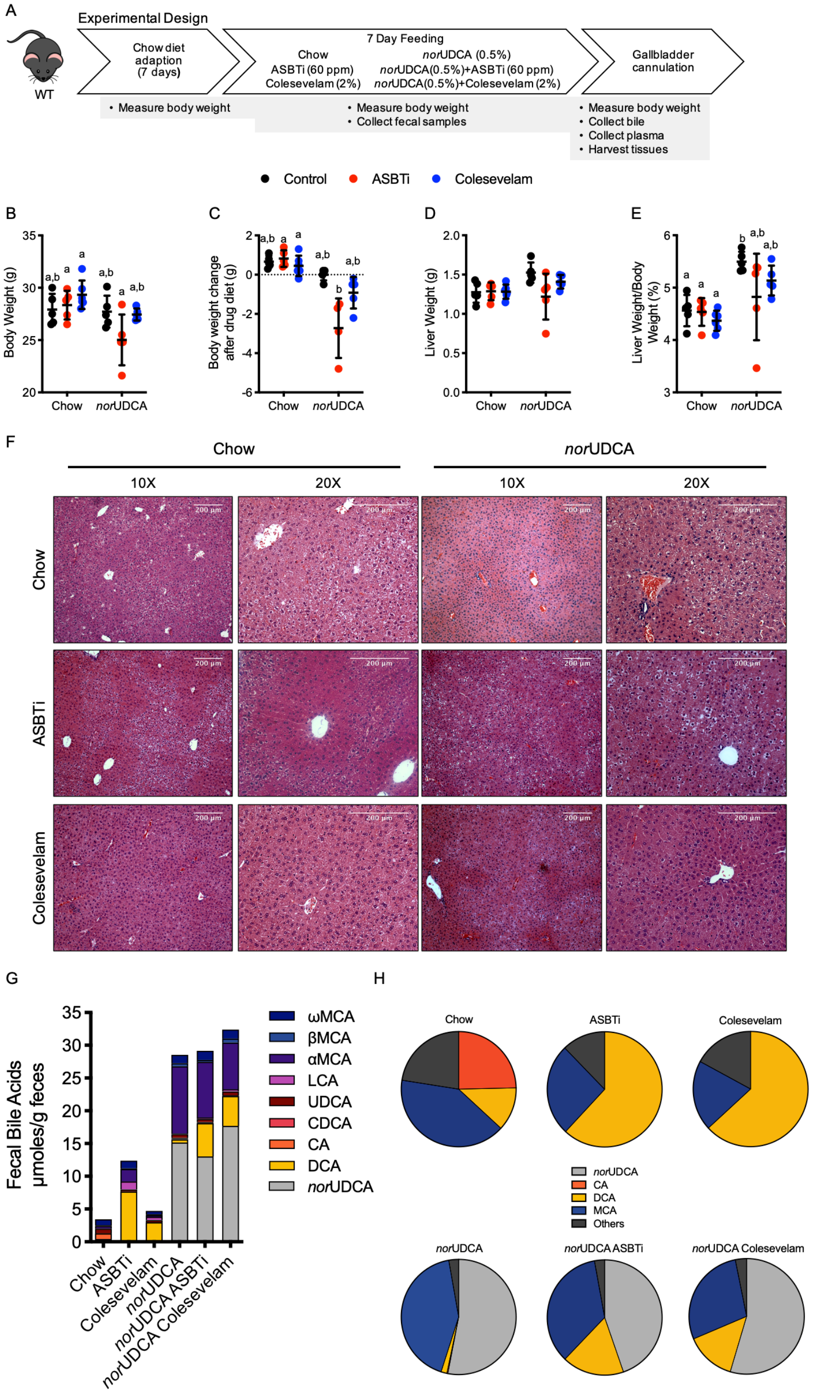


**Supplemental Figure 8. Experimental scheme and morphological response of WT mice to co-administration of *nor*UDCA with an ASBT inhibitor or Colesevelam.** (**A**) Experimental design. (**B**) Body weight. (**C**) Body weight change. (**D**) Liver weight. (**E**) Liver to body weight ratio. Mean ± SD, n = 5 mice per group. Distinct lowercase letters indicate significant differences between groups ($P < 0.05$). (**F**) Hematoxylin and eosin-stained liver sections (original magnification 10x or 20x) from the indicated treatments groups. *Scale bar*, 200 µm. (**G**) *nor*UDCA treatment increased the excretion of endogenous bile acids and the amount of 6-hydroxylated bile acids in feces. (**H**) Pie charts for the fecal bile acid profiles. Mean values are shown for fecal samples collected from cages of group-housed mice (n = 5 mice per treatment condition).

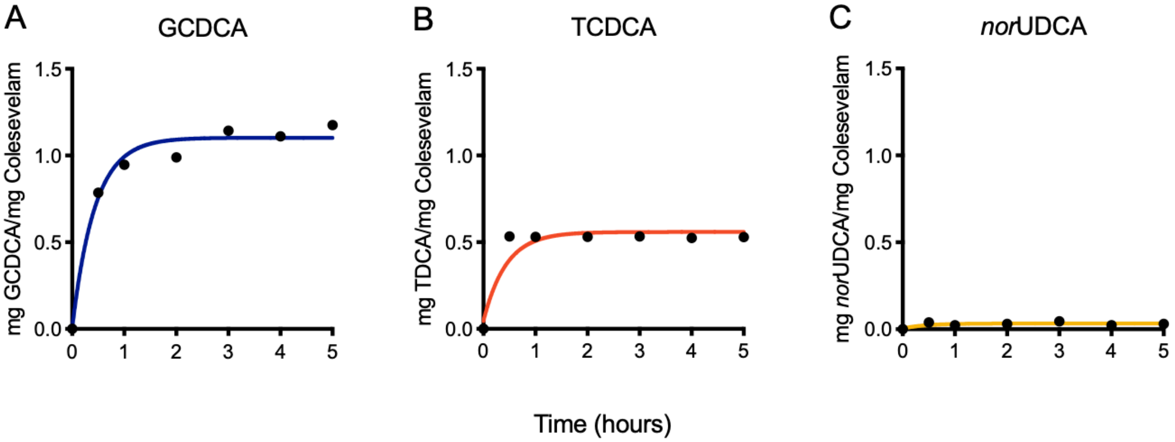


**Supplemental Figure 9. *In vitro* binding of bile acids and *nor*UDCA to colesevelam.** (**A**) Glycochenodeoxycholic acid (GCDCA), (**B**) Taurochenodeoxycholic acid TCDCA), (**C**) *nor*UDCA. Bile acids were dissolved in simulated intestinal fluid at a concentration of 2 mM for TCDCA or 6 mM for GCDA and *nor*UDCA and incubated with 7.5 mg of colesevelam hydrochloride at 37°C. Samples were withdrawn at 0, 0.5, 1, 2, 3, 4 and 5 h, filtered through a 0.2 μm syringe filter, and the filtrate was analyzed by enzymatic assay to determine the free bile acid concentration by enzymatic assay. The difference between the initial (0 h) and unbound bile acid concentrations at each time point was used to calculate the amount of the bile acid bound per mg of colesevelam hydrochloride.
